# Le click c’est chic: a plug-and-play virus-like particle vaccination platform enabled by non-canonical amino acid incorporation and click chemistry in the tobacco BY-2 cell-free protein synthesis system

**DOI:** 10.1101/2025.08.21.671528

**Authors:** Jorge Armero Gimenez, Janis Schleusner, Ruud Wilbers, Arjen Schots, Livia Spiga, John Tregoning, Ricarda Finnern, Charles Williams

## Abstract

Non-canonical amino acids (ncaas) can provide recombinant proteins with novel exciting functionalities beyond the limits of nature, such as orthogonal reaction groups. Notably, ncaa introduction can be used in vaccinology to enhance the adaptability and immunogenicity of putative vaccine candidates. Cell-free protein synthesis (CFPS) represents the most promising methodology to introduce ncaa into recombinant proteins of interest. However, traditionally used prokaryotic CFPS systems show limitations to produce complex proteins requiring post-translational modifications, whilst eukaryotic CFPS systems have historically been difficult to scale and show low protein yields. In this work, we establish the site-specific introduction of ncaas into complex proteins with the high-yielding and scalable eukaryotic tobacco BY-2 CFPS system (BYL), commercialised as ALiCE®. The tyrosine transferase from *Escherichia coli* (eTyrT) was tested for amber suppression-mediated ncaa incorporation in BYL. eTyrT showed high incorporation yields of up to 2mg/ml recombinant protein for the azido-tyrosine and alkyne-tyrosine ncaas, with linear scalability up to 10ml without any losses in protein yield. We applied ncaa incorporation in BYL to enable click chemistry bioconjugation of the receptor binding domain (RBD) of influenza hemagglutinin to pre-assembled hepatitis B core (HBc) virus-like particles (VLPs). BYL efficiently produced the alkyne-modified RBD and azido-modified HBc VLPs, and their conjugation via copper-catalysed azide-alkyne cycloaddition (CuAAC) led to structurally intact, RBD-coated particles. VLP-RBD conjugates could efficiently hemagglutinate chicken erythrocytes where the individual proteins could not, proving both the sialic-acid binding activity of the RBD and its multivalent presentation by the HBc VLP. Finally, when used to vaccinate mice the conjugated RBD-VLPs showed a greater protection against live influenza challenge than free RBD. This research thus enables ncaa introduction for recombinant proteins produced in BYL, constructing a novel plug-and-play vaccine platform and further expanding the capabilities of BYL to produce vaccine candidates and other proteins of interest.

## 1. Introduction

The production of recombinant proteins has become increasingly important due to their significant roles in medical and industrial applications (1). Subunit vaccines have gained growing interest following the global SARS-CoV-2 pandemic, which highlighted the need for rapidly adaptable vaccine platforms (2). Legacy protein-based vaccine modalities produced by traditional cell-based methods are limited in this context by their lengthy development times, thus limiting their applicability against emerging threats (3,4). Recent developments in the cell-free protein synthesis (CFPS) field suggests a faster and more adaptable alternative.

Instead of living cells, CFPS comprises cell extracts that contain the molecular machinery required for protein synthesis (transcription machinery, ribosomes, amino acids, tRNAs, etc.). CFPS thus only requires the addition of template DNA containing the sequence of interest to rapidly produce the protein (5,6). However, the majority of described CFPS systems are derived from prokaryotic *Escherichia coli,* showing limitations to produce complex proteins with post-translational modifications (PTMs), needed for human-relevant therapeutics and vaccines. Eukaryotic CFPS systems with PTM machinery have historically been difficult to scale and show low protein yields (7). The tobacco BY-2 cell-free lysate (BYL) has recently been developed as a novel and highly promising eukaryotic CFPS system, demonstrating higher yields of complex proteins than other CFPS systems with gram-scale protein production from 24- to 48-hour reactions in bioreactors (8,9). This led to its commercialization as ALiCE® by LenioBio GmbH.

The lack of cell walls and membranes in CFPS creates an open system that allows the straightforward modification of reaction conditions, thus permitting the addition of chemical and biological components that could be exploited for novel vaccine candidate development (10). This characteristic of CFPS has been used to introduce orthogonal biosynthesis pathways for the incorporation of non-canonical amino acids (ncaas) to recombinant proteins. Ncaas are synthetic structures that resemble amino acids, but that have been modified to provide specific functions to the protein in which they are incorporated. Such modifications include the addition of stabilizing groups, fluorescent dyes or specific reactive handles (11,12), the later allowing proteins to be covalently attached to other molecules (13–16). For vaccine development, ncaas have been used to increase the immunogenicity of the otherwise poorly immunogenic bacterial surface glycans: by conjugating the glycan via click chemistry to a carrier protein, an enhanced immune reaction could be achieved (17). Additionally, bioconjugation of subunit-type vaccines to virus-like particles (VLPs) via ncaas can exploit the adjuvant properties of the VLP via the multivalent presentation of a desired antigen, thus triggering a strong and specific immune response (18). This represents an advantage over antigen presentation in VLPs via protein fusion, as post-assembly bioconjugation does not limit antigen size, nor post-translational modifications of the epitope to be presented and allows for control over the number of antigen copies per VLP (19–21).

In this study, we advanced the BYL platform by enabling site-specific incorporation of ncaas with an amber suppression approach. An orthogonal tRNA/synthetase pair was introduced to code for the amber stop codon (UAG), introducing either azido-tyrosine or alkyne-tyrosine to proteins expressed in the BYL system (Figure 1). Ncaa-modified proteins are then suitable for bioconjugation through copper-catalysed azide alkyne cycloaddition (CuAAC) click chemistry. We applied this technology to construct a plug-and-play vaccine platform comprised of hepatitis B core antigen (HBc) carrier VLPs. These carrier VLPs can be conjugated to other recombinant antigens to boost their protective potential on vaccination, exemplified here with influenza antigens, and supported by *in vitro* and *in vivo* functional characterizations (Figure 1).

**Figure 1.**
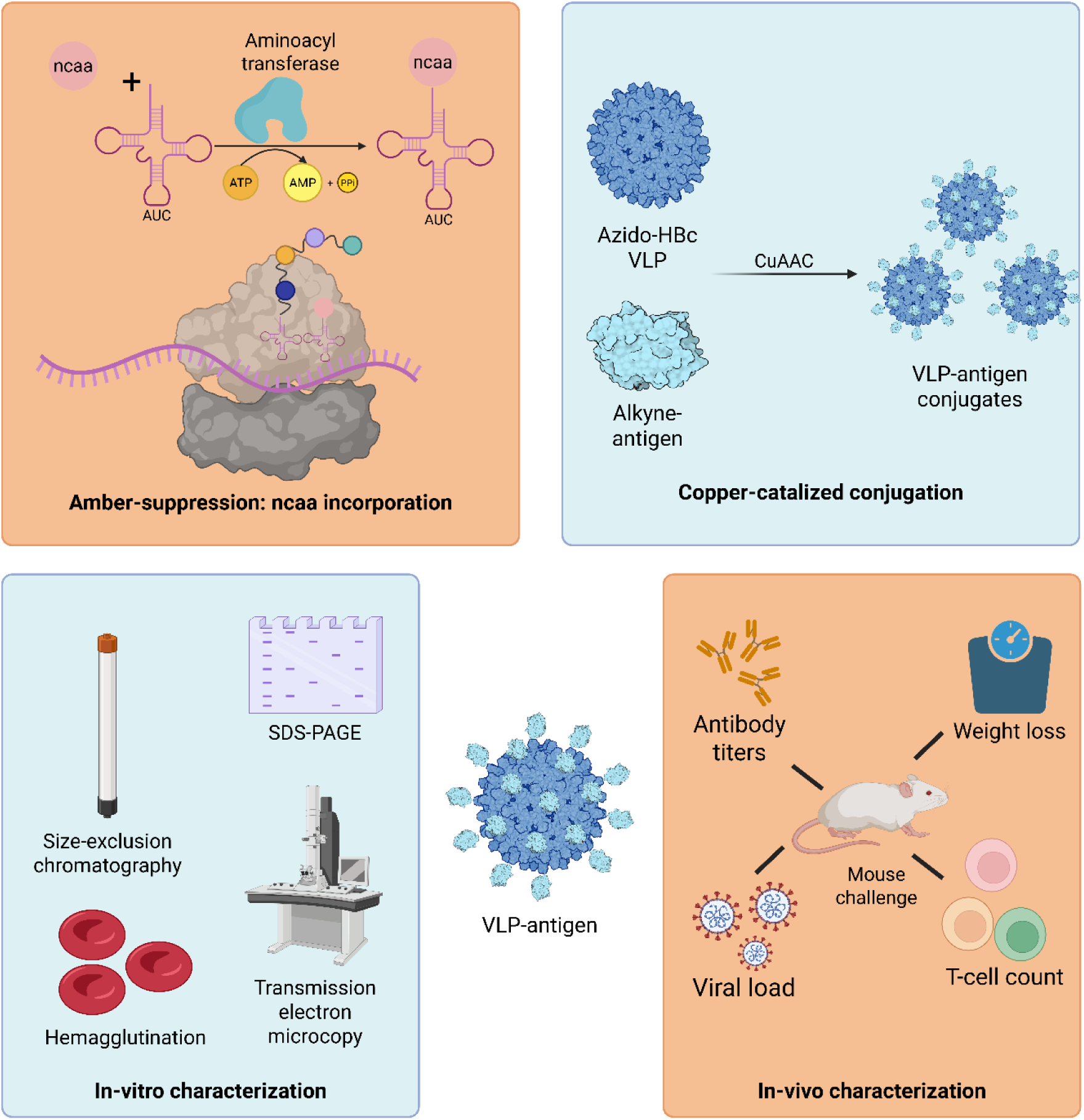
Ncaa incorporation in BYL for a plug-and-play vaccine platform. The aminoacyl-transferase specifically links the ncaa to the tRNA containing the AUC anti-codon. The ncaa-tRNA is used to code for the amber-codon, leading to the site-specific introduction of the ncaa into the nascent protein sequence. The azido-VLP and alkyne-antigen proteins produced in BYL can then be used in a copper catalysed click-reaction, resulting in their covalent conjugation. A series of *in vitro* and *in vivo* characterization techniques can then be performed to analyse obtained VLP-antigen conjugates. Abbreviations: BYL: tobacco BY-2-based cell-free protein synthesis system; ncaa: non-canonical amino acid; HBc VLP: hepatitis B core virus-like particle; CuAAC: Copper(I)-catalysed azide-alkyne cycloaddition. Created with BioRender.com.

We first tested and optimized the incorporation of ncaas in BYL with the tyrosine transferase from *E. coli* (eTyrT), and the pyrrolysine transferases from *Methanosarcina mazei* (mPyrT) and *Methanosarcina barkeri* (bPyrT) (22,23). Purified transferase supplementation to BYL was essential to reach high ncaa incorporation yields. The eTyrT system led to the site-specific incorporation of azido-tyrosine and alkyne-tyrosine into BYL-produced proteins, reaching yields between 90%-100% of the maximum BYL yield. The eTyrT also showed linear scalability up to a 10mL reaction volume with protein yields of >2 mg/mL. The functionality of the introduced alkyne and azido moieties was then proven by the production and bioconjugation of azido-modified HBc VLPs to alkyne-modified receptor binding domain (RBD) from influenza hemagglutinin. CuAAC reaction conditions could be optimised to control VLP conjugation efficiency by modulating the relative concentrations of the proteins. Characterisation of VLP-RBD conjugates by size-exclusion chromatography and transmission electron microscopy showed that structural integrity of the particles was maintained after the click chemistry reaction. A hemagglutination assay further demonstrated multivalent presentation of sialic acid-binding VLP-RBD conjugates. When used to immunize naïve mice, the VLP-RBD model showed enhanced protection against the development of disease when compared to free RBD, indicating the potential increase in immunogenicity upon conjugation to the HBc VLP carrier and validating the plug-and-play vaccine platform.

Taken together, we demonstrated the high-yield site-specific incorporation of ncaa into proteins produced in the BYL system and proved their activity in terms of their conjugation via click-chemistry. This research further expands the applicability of the BYL system to produce plug-and-play VLP-based vaccine candidates and other medically relevant ncaa-containing proteins.

## 2. Materials and Methods

### 2.1 Genetic construct design

The *M. mazei*, *M. barkeri* pyrrolysine and *E. coli* tyrosine tRNA sequences were retrieved from NCBI (GenBank: CP042908.1 & X69401.1) and synthesized by Integrated DNA Technologies (IDT) including the TAG anticodon. Note that the pyrrolysine tRNA sequence for *M. mazei* and *M. barkeri* is the same. A T7 promoter was used to drive the transcription of all tRNA transcripts. A hammerhead ribozyme sequence was included after the T7 promoter, consisting of the following sequence: NNNNCTGATGAGTCCGTGAGGACGAAACGGTACCCGGTACCGTC. The first 4 hammerhead nucleotides were switched to match the 4nt in the pyrrolysine (TTCC) and tyrosine (CACC) tRNA hybridization boxes (24,25). A version of these plasmids without the hammerhead ribozyme and a shorter T7 promoter were also constructed and tested. PCR templates obtained from these plasmids were utilized for *in vitro* transcription (IVT) and BYL tRNA co-transcription.

The protein sequences for the *M. mazei*, *M. barkeri* and *E. coli* aminoacyl-transferases were retrieved from NCBI (GenBank: WP_011033391.1, ACB70968.1 & ALJ52437.1, respectively), codon optimized for expression in *Nicotiana tabacum* and synthesized by IDT. Gibson cloning was used to clone the engineered coding sequence into the pALiCE01 expression plasmid as previously described (26). Expression plasmids were purified from *E. coli* DH5alpha cultures using the NucleoBond Xtra Maxi kit (Macherey Nagel) and their correct assembly was confirmed by sequencing (Eurofins). A reporter construct based on enhanced yellow fluorescent protein (a-eYFP) was designed by introducing an amber codon in a loop structure before the fluorophore at amino-acid position 50 of the eYFP (69 of the pALiCE01 Strep-tagged eYFP construct). Thus, a quantifiable fluorescent response is only observed upon correct introduction of the ncaa and complete protein translation.

A dimer fusion of the HBc VLP monomer referred to as “tandem core” was used for VLP production (27). A C-terminal His-tag was included for purification. Linkers of different length were included into the MIR position of HBc VLP, resulting either in a no-linker construct (tyr-VLP and az-VLP), one with a small GSSG*GSSG linker (tyr-short and az-short), and another with a long GGGGSGGGG*GGGGS GGGG linker (tyr-long and az-long). The * represents either a tyrosine site or an amber codon site, thus resulting in 6 different HBc VLP tandem core constructs. The influenza hemagglutinin RBD sequence (amino acids 63 to 286 (28)) from influenza A virus (A/Puerto Rico/8-SV1/1934) was retrieved from NCBI (GenBank: ACO94826.1), codon optimized for expression in *N. tabacum* and synthesized by IDT. The coding sequence was cloned into pALiCE01 for cytosolic BYL expression. Four different RBD-expressing constructs were obtained, either containing an N- or C-terminal Strep-tag for purification, and a tyrosine or amber codon site. The tag contained an HRV-3C protease recognition signal prior linker sequences, resulting in the sequences: MAWSHPQFEKGGSLE VLFQGPGGS*GGSGGSG & GGSGGS*GGSLEVLFQGPGG SAWSHPQFEK, for N-terminal and C-terminal tags, respectively. The * indicates either the amber stop codon or a tyrosine residue for the non-canonical and canonical RBD constructs, respectively.

### 2.2 tRNA template production

The primers required to generate the PCR templates for tRNA transcription included 4 phosphorothioate (PTO) linkages to prevent exonuclease degradation in BYL. The forward primer was designed to bind ∼400bp upstream the T7 promoter. The reverse primer also included a methoxy modification in the second to last nucleotide to prevent untemplated addition at 3’ by the T7 polymerase (29,30). PCR reactions to generate the tRNA templates were performed using Phusion™ High-Fidelity DNA Polymerase (New England Biolabs) with the GC buffer and following instructions provided by the manufacturer. PCR purification was performed using the NucleoSpin Gel and PCR Clean-up kit (Macherey-Nagel). To use oligos as template, Ultramer primers (IDT) were ordered. For the forward strand, either a short primer encompassing only the T7 promoter, or an ultramer primer covering the whole tRNA sequence were ordered, both containing 4-PTO linkages in the first 5nt. The reverse strand primer covered the whole T7 + tRNA cassette, containing the same PTO linkages and the methoxy modification in the second to last nucleotide as previously indicated. Oligos were denatured at 95°C for 5min, and then annealed by slowly cooling to 25°C. To use linearized plasmids tRNA template, a BsaI restriction site was placed downstream of the tRNA CCA acceptor arm, allowing run-off transcription. Digestion with BsaI (New England Biolabs) was performed as per manufacturer’s instructions, followed by inactivation at 65°C for 20 min, and purification using the NucleoSpin Gel and PCR Clean-up kit (Macherey-Nagel) as previously described.

### 2.3 *In vitro* transcription and analysis

*In vitro* transcription (IVT) reactions were performed using the HiScribe T7 Quick High Yield RNA (New England Biolabs) as per manual instructions, using 1 µg of purified DNA as template. To test the efficiency of the hammerhead ribozyme, a 1 h heat treatment at 60°C was performed, either without or with the addition of 2 mM MgCl_2_. After reaction completion, 2x RNA dye (New England Biolabs) was added, and samples were incubated at 60°C for 10 min and immediately cooled on ice prior to loading. Samples were run in 0.5x TB buffer (45 mM Tris, 45 mM boric acid) at 250 V for 20 min in 4% Agarose gels as previously described (31), SYBR gold staining (ThermoFisher Scientific) was used as a dye for gel visualization. Pictures were taken using a Gel Doc XR+ UV transilluminator (Bio-Rad). Alternatively, *in vitro*-transcribed tRNAs were analysed via TBE-Urea PAGE using precast Novex™ TBE-Urea Gels, 10% (ThermoFisher Scientific). Samples were denatured in 2x Novex TBE-Urea Sample Buffer (ThermoFisher Scientific) and run for 60 min at 180 V in 1X Novex TBE Running Buffer (ThermoFisher Scientific). Gel staining and detection was performed as previously stated. tRNA quantification after IVT was performed using the QuantiFluor RNA System (Promega).

### 2.4 BYL protein production, purification and analysis

BYL expression was performed as previously described (32). Briefly, 50 and 100 µl BYL reactions were set by thawing the BYL lysate in a water bath at room temperature, and each expression plasmid was added to reach 5 nM. Reactions were incubated at 25°C with 75% humidity in half-well and full-well 96-well plates for 50µl and 100µl reactions, respectively, at 500 rpm with a 12.5 mm shaking diameter. Alternatively, 5 ml and 10 ml scale BYL reactions for purification were run in a LT-X or ISF1-X Kuhner shaker for 48 h in sterile 100 ml and 250 ml Erlenmeyer flasks, respectively, at 25°C and 95 rpm with a shaking diameter of 50 mm. After reaction completion, resulting lysate was centrifuged at 16,000 x *g* for 10 min to remove insoluble debris, and soluble recombinant transferases were purified via immobilized-metal affinity chromatography (IMAC) using PureCube 100 Compact Cartridge Ni-NTA (Cube Biotech) prepacked columns using an Äkta chromatography device (Cytiva). For HBc VLP purification, IMAC was performed under denaturing conditions, and re-folding was induced after purification by overnight dialysis into re-assembly buffer (50 mM Tris HCl pH 7, 800 mM NaCl). Purification of RBD was performed Strep-Tactin®XT 4Flow® high capacity FPLC column (IBA), also under denaturing conditions, with an in-column re-folding step as previously described (33). .

Protein samples were analysed using NuPAGE™ 4 to 12%, Bis-Tris (Invitrogen™) SDS-PAGE pre-cast gels. Protein samples were prepared according to the manufacturer’s instructions. 5μl of PageRuler™ Prestained Protein Ladder (ThermoFisher Scientific) were also loaded as size standard. Gels were then run in 1x MES running buffer (ThermoFisher Scientific) for 25min at 200V or until sufficient resolution was obtained. Successively, gels were stained with Coomassie Blue-staining solution (25% (v/v) isopropanol, 10% (v/v) acetic acid and 0.05% (w/v) Coomassie brilliant blue R-250) and destained with 10% acetic acid until protein bands were clearly discernible. Gel images were captured using the Gel Doc XR+ UV transilluminator (Bio-Rad Laboratories, Inc.).

### 2.5 Non-canonical amino-acid incorporation in BYL

Ncaa incorporation via co-expression was performed via the supplementation of reactions expressing the transferase and the tRNA with their respective ncaa after 8 or 24h, where different % (volume initial reaction)/(volume final reaction) was transferred to a new BYL reaction expressing a-eYFP. The ncaa used in this research were H-L-Phe(4-N3)-OH (azido-tyrosine) and H-L-Tyr (Propargyl)-OH (alkyne-tyrosine), for the eTyrT, and H-L-Lys (EO-N3)- OH*HCl (azido-pyrrolysine) for the pyrrolysine transferases. All three ncaa chemicals were purchased from Iris Biotech, with 100mM stock solutions prepared in 20% DMSO and were included in the BYL reaction at a 5 mM final concentration. Fluorescence signal after reaction completion was measured using an Infinite M1000 device (Tecan Group Ltd.), with excitation and emission wavelengths of 485 nm and 528 nm, respectively. For experiments where the transferase was externally produced, purified, and supplemented, BYL reactions were set as previously described and tRNA PCR template was included at concentrations ranging from 25nM to 75nM and the transferases were tested in concentrations ranging from 1µM to 5 µM.

### 2.6 CuAAC reaction

Copper(I)-catalysed azide-alkyne cycloaddition (CuAAC) reactions were performed as previously described (34). Briefly, 3-azido-7-hydroxicoumarin (Baseclick) substrate was used to determine the optimal CuAAC reaction conditions in presence of HBc VLP, by measuring fluorescence after reaction with 10 µM propargyl alcohol, with an excitation and emission wavelengths of 404 nm and 477 nm, respectively. HBc VLP to RBD conjugation reactions were performed by combining each of the coupling partners in a reaction vessel as to reach the indicated molar ratios alkyne-RBD to azide-VLP with a minimum protein concentration of 10 µM for the Azido-VLP. Final CuAAC reaction conditions were 250 µM CuSO_4_, 1.25 mM THPTA ligand, 5 mM Na-Ascorbate, 500 µM NiCl2 and 5 mM aminoguanidine. Reactions were incubated for 1 hour at room temperature, after which EDTA was added as to reach 10 mM to stop the reaction. Reaction yields were then estimated via SDS-PAGE electrophoresis and gel densitometry analysis using GelAnalyzer software.

### 2.7 SEC and TEM analysis

Resulting RBD-VLP conjugates were analysed by size exclusion chromatography (SEC) and transmission electron microscopy (TEM). SEC analysis was performed using a Superdex 200 column (Cytiva) on an ÄKTA purifier and was also used as a purification step to remove unbound alkyne-RBD from CuAAC coupling reactions prior to further testing. For TEM analysis, formvar-carbon coated 400 mesh copper grids (Sigma-Aldrich) were glow discharged in vacuum for 20 s. 10 μl of each VLP suspension were placed on a grid and incubated for 2 min. Negative staining was performed with 1% (v/v) phosphotungstic acid (PTA, pH 7.2) for 1 min. The specimens were examined in a JEOL 1400 transmission EM equipped with a Matataki (2K × 2K).

### 2.8 Hemagglutination assay

Chicken erythrocytes were purchased from Fiebig GmbH and washed three time with PBS supplemented with 2 mM Penicillin/Streptomycin. 50µl samples were plated in 96-well round well plates at the indicated concentrations, and a 1:1 dilution series was performed column-wise. Subsequently, 50µl of 1% (v/v) erythrocytes were added to each well, and the plate was incubated for 1 h at 4°C. Hemagglutination titre could then be measured by the last serial dilution that induces erythrocyte agglutination. Commercial inactivated influenza vaccine (kindly provided by Herr Dr. Keul), and hemagglutinin protein (Abcam) were used as controls.

### 2.9 Mouse immunisation

For experiments to test protection against infection, 6–10 week old female BALB/c mice were obtained from Harlan UK Ltd (Liverpool, UK) and kept in specific-pathogen-free housing. Studies followed the ARRIVE guidelines. Mice were immunised with one of the following formulations: free RBD, RBD conjugated to VLP, free RBD combined with free VLP, and free VLP as negative control. Mice were immunised intramuscularly with 5 μg of the recombinant vaccine formulated with a 1:1 dilution with AddaVax adjuvant (InvivoGen) in 50 μl in a prime-boost-boost regimen. For infections, mice were anesthetized using isoflurane and infected intranasally with 100 μl influenza virus or sterile PBS. Mice were culled using 100 μl intraperitoneal pentobarbitone (20 mg dose, Pentoject, Animalcare Ltd.) and tissues collected as previously described (35). Viruses were propagated in Madin-Darby Canine Kidney (MDCK) cells, in serum-free DMEM supplemented with 1 µg/ml trypsin. Influenza viral load was assessed by PCR as described previously (36).

### 2.10 ELISA assay

HA-specific serum IgG were evaluated by enzyme-linked immunosorbent assay (ELISA) as previously described (37). Briefly, MaxiSorp microtitre plates (Nunc) were coated with recombinant PR/8 HA (1 µg/ml: Sino) overnight at 4°C, blocked with PBS and 1% BSA, and then added with serum samples titrated in two-fold dilutions. Samples were then incubated with the alkaline phosphatase-conjugate goat anti-mouse IgG (diluted 1:1000, Southern Biotechnology) for 1 hour at 37°C and developed by adding 1 mg/ml of alkaline phosphatase substrate (Sigma-Aldrich). The optical density was recorded using Multiskan FC Microplate Photometer (Thermo Scientific).

### 2.11 Viral Load by RT-PCR

Lung tissue was homogenized in Trizol using the TissueLyzer (Qiagen) instrument at 50 oscillation for 4 min then RNA was extracted from homogenates using TRIzol/chloroform extraction. Briefly, homogenates were transferred to 1.5 mL tubes, and 150µL TRIzol reagent was added, followed by a 5 min RT incubation. 100µL chloroform was then added, and the tubes were inverted to mix. After a 15 min centrifugation at 13,000 x *g* at 4 °C, the aqueous phase was transferred to fresh tubes. RNA was precipitated with 250µL isopropanol (10 min RT incubation and 10 min centrifugation at 13,000 x *g*), washed with 75% ethanol (5 min at 7,500 x *g*), air-dried for 15 min, and resuspended in nuclease-free H2O. Samples were incubated at 55 °C for 5 minutes to denature RNA secondary structures and then placed on ice. Total RNA concentrations and purity were determined using a NanoDrop One (ThermoFisher Scientific) spectrophotometer. All RNA samples were normalised to 200ng/µL and converted into cDNA using a GoScript reverse transcription system (Promega). qRT-PCR was performed for the influenza M gene on a Stratagene Mx 3005p (Agilent Technologies) instrument, using primers 5′- AAGACAAGACCAATYCTGTCACCTCT-3′ and 5′-TCTACGYTGCAGTCCYCGCT -3′ and probe 5′-FAM-TYACGCTCACCGTGCCCAGTG-TAMRA-3. M gene-specific RNA copy number was determined against an Influenza M gene standard.

Statistical analysis was performed using GraphPad Prism. A one-way ANOVA with multiple pairwise comparisons was used to compare group means, applying Dunnett’s post hoc test to adjust for multiple comparisons. A p-value of <0.05 was considered statistically significant.

## 3. Results

### 3.1 Characterisation of amber suppression components

In this study, we investigated the site-specific ncaa incorporation in the BYL CFPS system using the amber suppression systems from *E. coli, M. mazei* and *M. barkeri* (eTyrT, mPyrT and bPyrT, respectively). For this purpose, the BYL translation machinery needs to be combined with the amber suppressor tRNA, along with their respective aminoacyl-transferase and ncaa. Firstly, we investigated tRNA transcription, for which different constructs with and without a 5’ hammerhead ribozyme were designed (Figure S1A). Agarose-gel electrophoresis after *in vitro* transcription (IVT) confirmed formation of the respective tRNA transcripts for both types of constructs, though addition of the hammerhead domain showed decreased transcript yields of the expected size. We subsequently tested expression of different aminoacyl transferase constructs in BYL (Figure S2A). BYL expression of the transferases led to considerably high yields (ca. 1mg/ml for eTyrT), observed as a visible band in Coomassie-stained SDS-PAGE gels (Figure S2B). However, the pyrrolysine transferases, especially bPyrT, showed lower yields of soluble protein, with some of it being present in the non-soluble fraction, even when the Smbp solubilization tag was utilized. Consequently, the eTyrT and mPyrT systems were chosen for further testing.

### 3.2 Establishing high-yielding amber suppression in BYL

Once the successful transcription and expression of each of the components required for amber suppression was confirmed, their use for ncaa incorporation in BYL could be assessed. Our initial approach for ncaa incorporation in BYL consisted of the expression of the transferase and tRNA constructs in BYL in presence of their respective ncaa (Figure S3). After the indicated reaction time (8 h or 24 h), a volume of this reaction was supplemented into a fresh BYL reaction expressing the amber-containing eYFP construct (a-eYFP), thus allowing the estimation of ncaa incorporation yield (Figure S3A). A great reduction on eYFP yield was observed when the supplementation was performed, with an even greater reduction when higher supplementation volumes and incubation times were used (Figure S3 B,D). This effect was more prominent for the eTyrT than for the mPyrT. Regarding ncaa incorporation, the eTyrT system showed higher yields than the mPyrT system, reaching 2% and 0.2% of the maximum BYL yield, respectively (Figure S3 C,E). Supplementation after 32h and 48h led to considerably lower ncaa incorporation yields (data not shown). Interestingly, similar incorporation yields were observed for the eTyrT when supplementing at 50 and 25% v/v, which could indicate resource competition of higher transferase yields leading to more efficient ncaa incorporation but lower a-eYFP expression. Consequently, we hypothesized that the external production and supplementation of the transferase to BYL could lead to greater ncaa incorporation yields.

Expression and purification of eTyrT and smPyrT for BYL supplementation was performed via IMAC. eTyrT was successfully purified whilst smPyrT showed the co-purification of other proteins or fragments (Figure S4). Purified aminoacyl-transferases and tRNA templates (containing the hammerhead sequence) were supplemented in fresh BYL reactions to drive ncaa incorporation (Figure 2, A-C). The eTyrT system efficiently incorporated both, azido- and alkyne-tyrosine, with yields of 90% to 100% of the maximum BYL yield for eYFP (Figure 2D). Ncaa incorporation using the tRNA template without a hammerhead led to lower incorporation yields (data not shown). Interestingly, azido-tyrosine incorporation was observed when no eTyrT was included, as shown in the orthogonality control 2. This could be indicative of partial incorporation of the azido-tyrosine into the tRNA by native transferases. No a-eYFP expression was observed when both the enzyme and the tRNA were included in absence of the azido-tyrosine or alkyne-tyrosine ncaa. This indicates that no native amino acid is being incorporated in the amber site. Very low ncaa incorporation yields were observed for the mPyrT system, with a maximum of just 1% of the maximum BYL eYFP yield when highest tRNA template concentrations were used (Figure S5). Utilizing the hammerhead-less construct led to a higher incorporation yield of ca. 6%, which could indicate that tRNA production and/or processing might be a limiting factor of mPyrT-mediated ncaa incorporation. Overall, we observed that the supplementation of pure aminoacyl-transferase was essential to reach considerable ncaa-incorporation yields in BYL, with the eTyrT system showing best results and thus being chosen as the ncaa incorporation system to be carried forward for further development in the following experiments.

**Figure 2.**
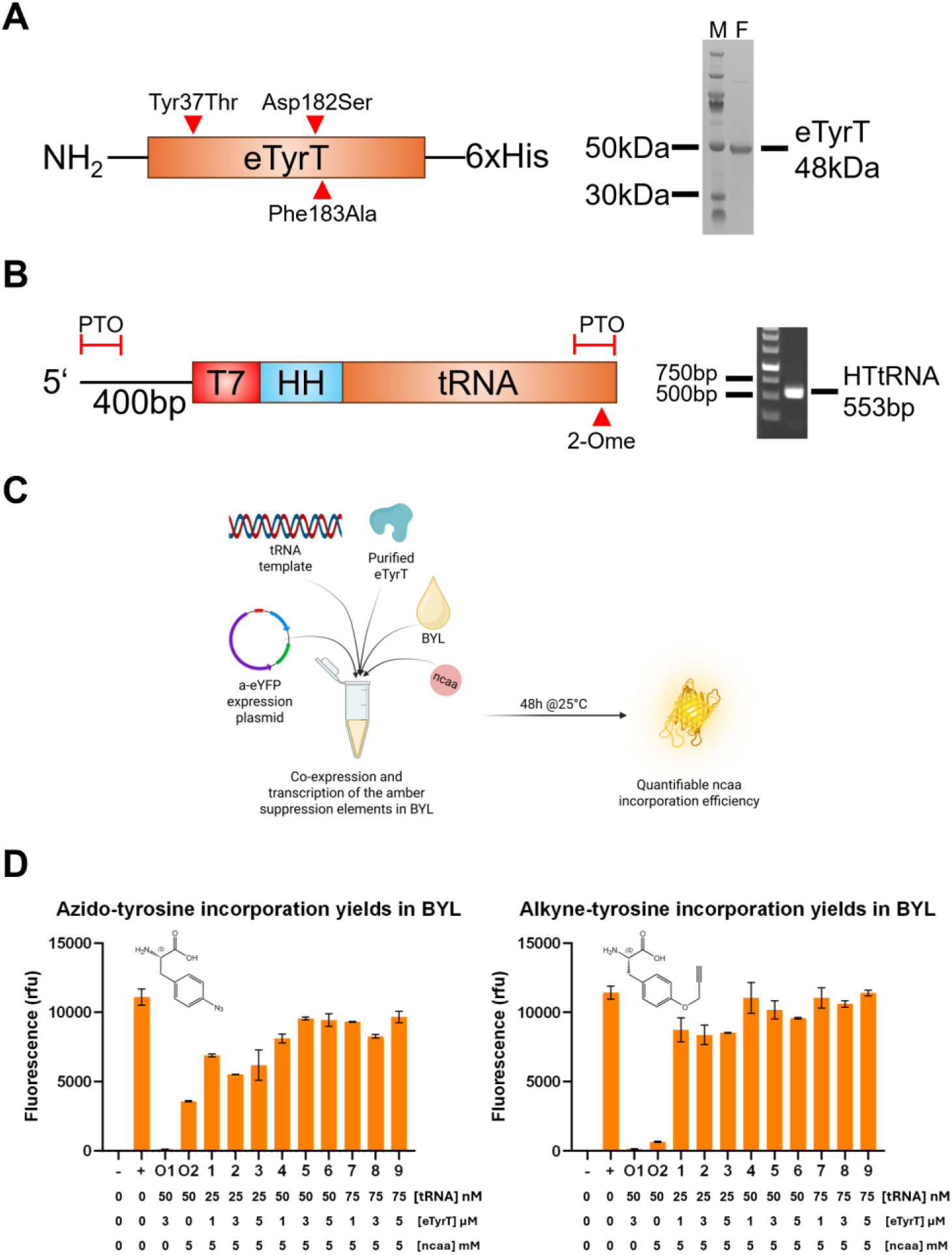
Enabling high-yield ncaa-incorporation in BYL. The *E. coli* tyrosine transferase amber suppression system was utilized to drive ncaa incorporation in the BYL cell-free protein synthesis system. For that purpose, the different components were first obtained. (A) Aminoacyl-transferase was expressed in BYL, containing the indicated mutations to enable the introduction of the ncaas, and purified. (B) The tRNA template used for the co-transcription of the tRNA was generated via PCR, including 4xPTO linkages at both ends, and a 2-Omethoxy modification in the second to last nucleotide. (C) Diagram of how the different components were combined to drive ncaa incorporation in BYL. (D) The incorporation efficiency was measured in terms of fluorescence of the amber eYFP construct after a 48h BYL reaction. Expression of eYFP is used as a positive control (indicated as +), indicating the maximum BYL yield, and the orthogonal controls 1 and 2 indicate the nonspecific incorporation of native amino acids and ncaa by native transferases, respectively. Different concentrations of the tRNA template and eTyrT enzyme were explored as indicated. Data points represent averages of experiments performed in duplicate. Abbreviations: eTyrT: *E. coli* tyrosine aminoacyl transferase; HTtRNA: Hammerhead tyrosine tRNA; rfu: relative fluorescence units; (-): Non-template BYL control; (+) pALiCE01 positive control; O1-2: orthogonality controls.

### 3.3 Scaling of amber suppression-enabled BYL reactions

An essential consideration in scalable protein production is maintaining high protein yields while ensuring the sustainable supply of additional reaction components. Production of eTyrT in scalable BYL reactions is well understood (8,38), but the custom tRNA PCR template needed to be addressed. Polymerase-chain reactions are not easily scalable, given limitations in heat transfer and the costs of its components (39,40). Alternative tRNA-generation approaches were thus devised to enable the scaling of ncaa incorporation in BYL. We explored the utilization of different DNA templates to obtain the tRNA, either directly within the ALiCE reaction or after IVT (Figure 3A). Annealing of short and long oligonucleotides was tested to support more cost-efficient co-transcription of the tRNA. Plasmid template linearised by restriction enzyme digest was also attempted to drive the run-off transcription of the tRNAs. All alternative constructs were tested with and without the hammerhead ribozyme. Co-transcription in BYL with short oligonucleotides showed no relevant a-eYFP production, whilst only ca. 60% of the PCR template yields were achieved when high concentrations of long oligonucleotides were used, and no transferase was added (Figure S6).

**Figure 3.**
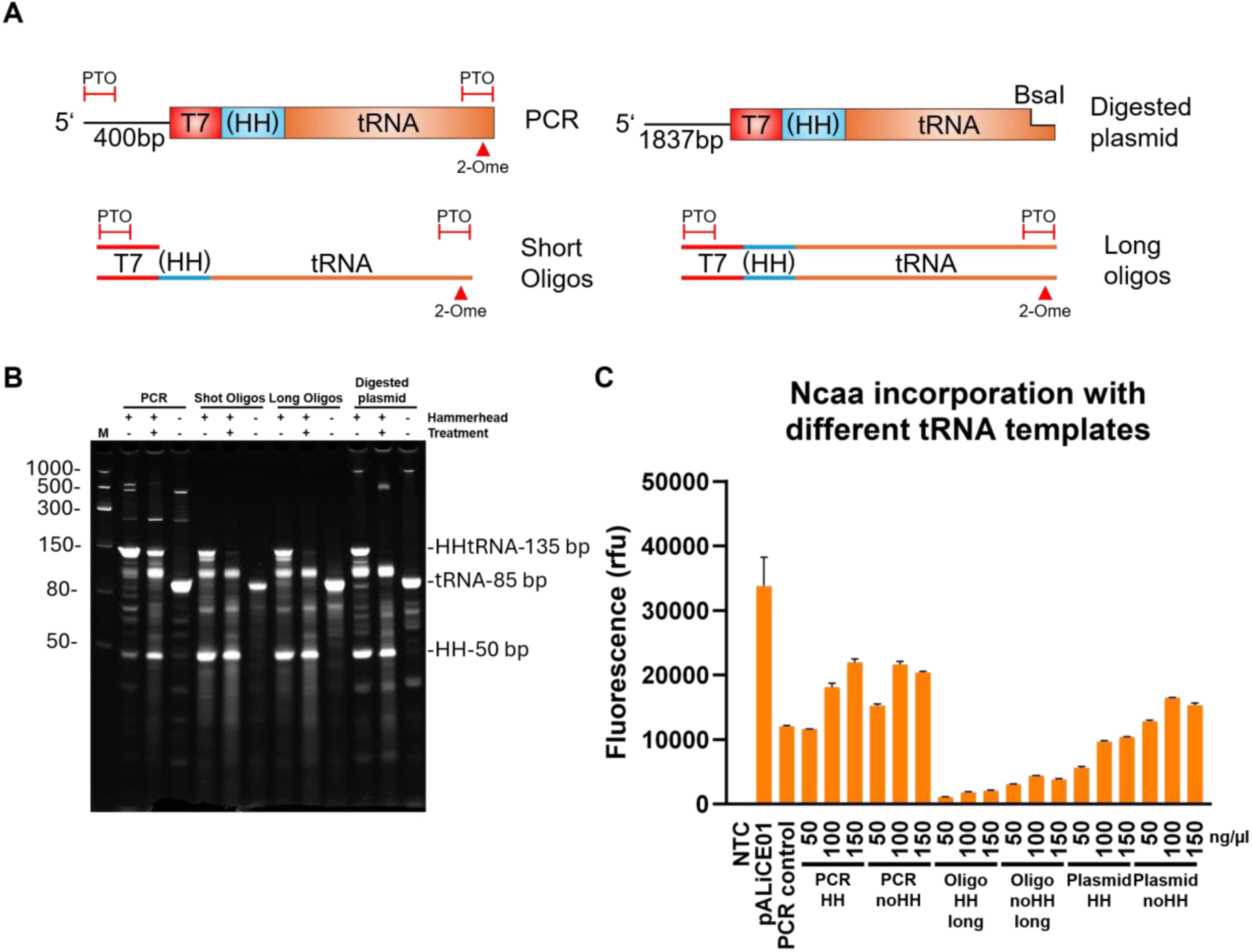
Alternative tRNA templates and *in vitro* transcription for ncaa incorporation in BYL. **(A)** Different tRNA templates and constructs were tested to enable ncaa incorporation at scale in BYL. Each tRNA template was generated with and without a 5’ hammerhead ribozyme to enhance 5’ homogeneity. **(B)** PAGE-Urea analysis of the *in vitro* transcription results from each construct and tRNA template. An 1h 60°C treatment was applied to constructs containing a hammerhead to assess cleaving activity. **(C)** BYL ncaa incorporation yields when utilizing the IVT products from each tRNA template at the indicated concentrations (ng/µl). The results from the short oligo are not shown as yields were negligible. No aminoacyl transferase was added in this experiment, instead relying on latent host transferase activity for this template screening assay. Data points represent averages of experiments performed in duplicate. Abbreviations: HH: hammerhead ribozyme; PTO: phosphorothioate linkage; 2-Ome: 2-O-methoxy-modification; M: molecular size marker; NTC: non-template control.

Uncoupled tRNA transcription was then tested to maximize ncaa incorporation yields. When transcribed *in vitro*, the different templates showed different behaviours depending on the presence of the hammerhead ribozyme (Figure 3B). Hammerhead-containing constructs seemed to generate a transcript which was longer than the expected 85bp tRNA. Shorter transcripts were also observed, indicating alternative ribozyme cleavages. When utilized in BYL, the different templates showed very different yields in ncaa incorporation: tRNA originating from PCR templates continued to show the highest yield, followed by linearised plasmid and the long oligonucleotide approach. Constructs with hammerhead domains neither increased nor reduced ncaa incorporation yields noticeably. IVT of tRNA from PCR template was thus progressed due to providing the most efficient ncaa incorporation, whereby the IVT process helps to amplify the tRNA amounts and support larger scale BYL reactions.

Subsequently, the scalability of this optimised approach was tested by performing 3 independent expressions at 4 different scales: 50µl, 100µl, 5ml and 10ml, the latter representing 200-fold increase from the standard reaction volume (Figure 4A). Ncaa incorporation in BYL showed linear scaling, with a yield of ca. 2mg/ml of recombinant protein across reaction volumes (Figure 4, B-C), thus enabling the introduction of ncaa in BYL at high yields and at scale.

**Figure 4.**
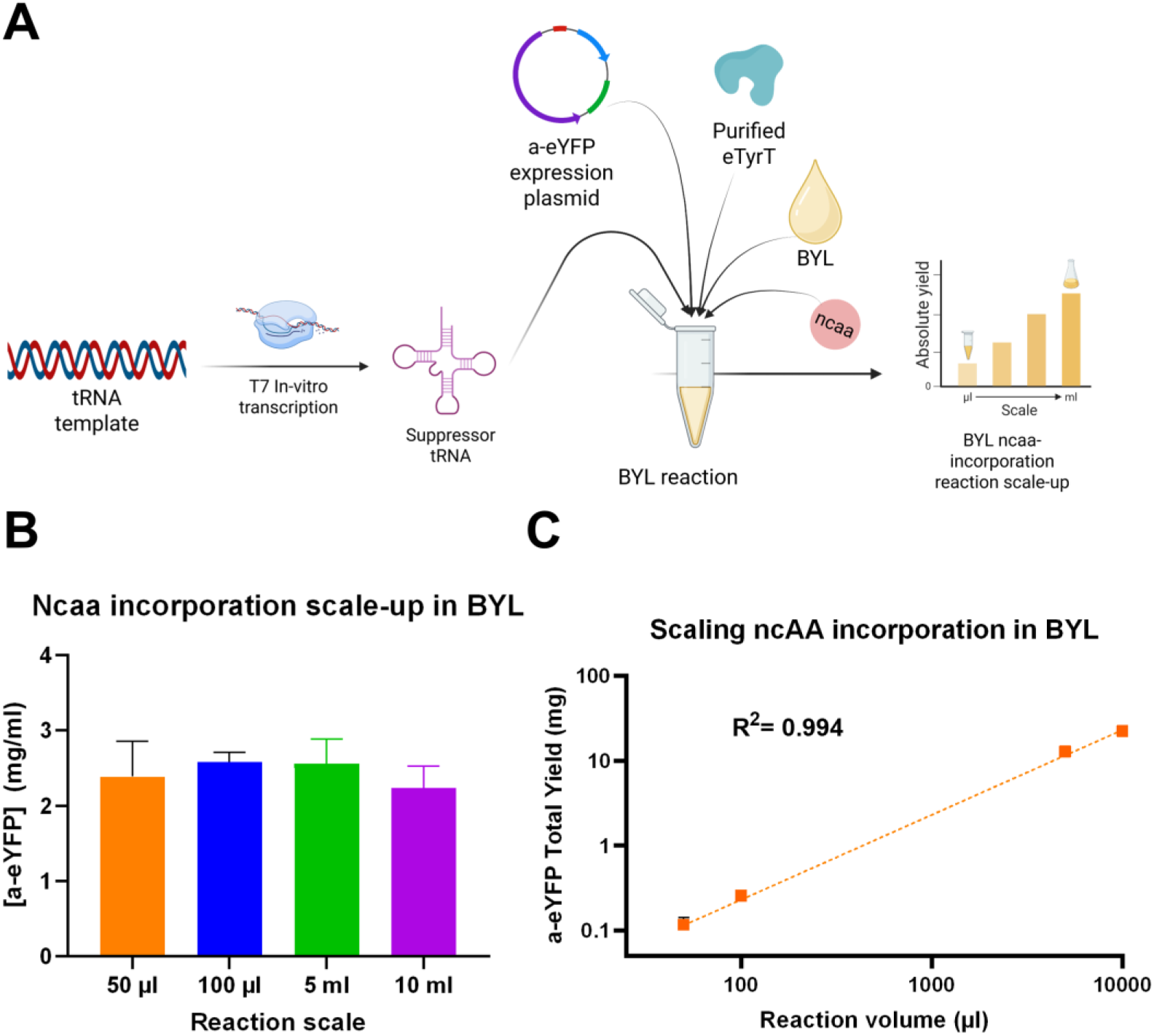
Ncaa incorporation scales linearly in BYL across a 200x scale range. (A) tRNA transcription was uncoupled from the BYL CFPS reaction to enable ncaa incorporation scale-up. *In vitro* transcribed tRNA was quantified and fed into a BYL reaction without purification to enable ncaa incorporation. (B-C) a-eYFP yields across the different tested scales, plotted as a-eYFP concentration (B) and total a-eYFP yield (C). Data points represent averages of experiments performed in triplicate. Abbreviations: a-eYFP: amber-eYFP; eTyrT: *E. coli* tyrosine aminoacyl transferase; BYL: tobacco BY-2 cell-free protein synthesis system.

### 3.4 Design, production and selection of amber suppression-compatible VLP and RBD constructs in BYL

We aimed to apply the newly developed amber suppression capabilities of BYL in a vaccinology context, where incorporation of ncaas to proteins could allow for click chemistry enabled enhancements. Specifically, a novel plug-and-play vaccine platform was envisioned, where hepatitis B core antigen (HBc) carrier virus-like particles (VLPs) containing azido-tyrosine (az-VLP) would be conjugated to subunit vaccine antigens via click chemistry. The production of functional HBc VLPs in BYL up to a 1L reaction scale was previously described (38). Here, additional tandem core HBc VLP constructs were designed to include either a tyrosine or ncaa insert at position 79 in the outwardly accessible major immunomodulatory region (MIR) of the HBc particle (Figure 5A). Two linkers of increasing length flanking the introduced tyrosine (tyr-VLP) or amber site (az-VLP) were also tested to determine whether any variant would present higher stability, assembly or conjugation yields. Expression of VLP constructs containing the Tyr residue in BYL showed high yields of recombinant protein, which could be efficiently purified by denaturing IMAC (Figure 5, B-D). However, constructs containing the linkers showed protein aggregation upon re-folding, with an overall loss of 60% to 85% for the longer and shorter linker constructs, respectively. No loss was observed for the VLP construct without linker. Therefore, the construct without a linker was chosen for ncaa incorporation. The production and purification of azido HBc VLP in BYL is shown in Figure 5E. Relatively high yields of over 80% of the tyr-VLP construct were obtained. VLP dimers promptly refolded into VLPs, independently of the incorporation of azido-tyrosine (Figure 5 F-G).

**Figure 5.**
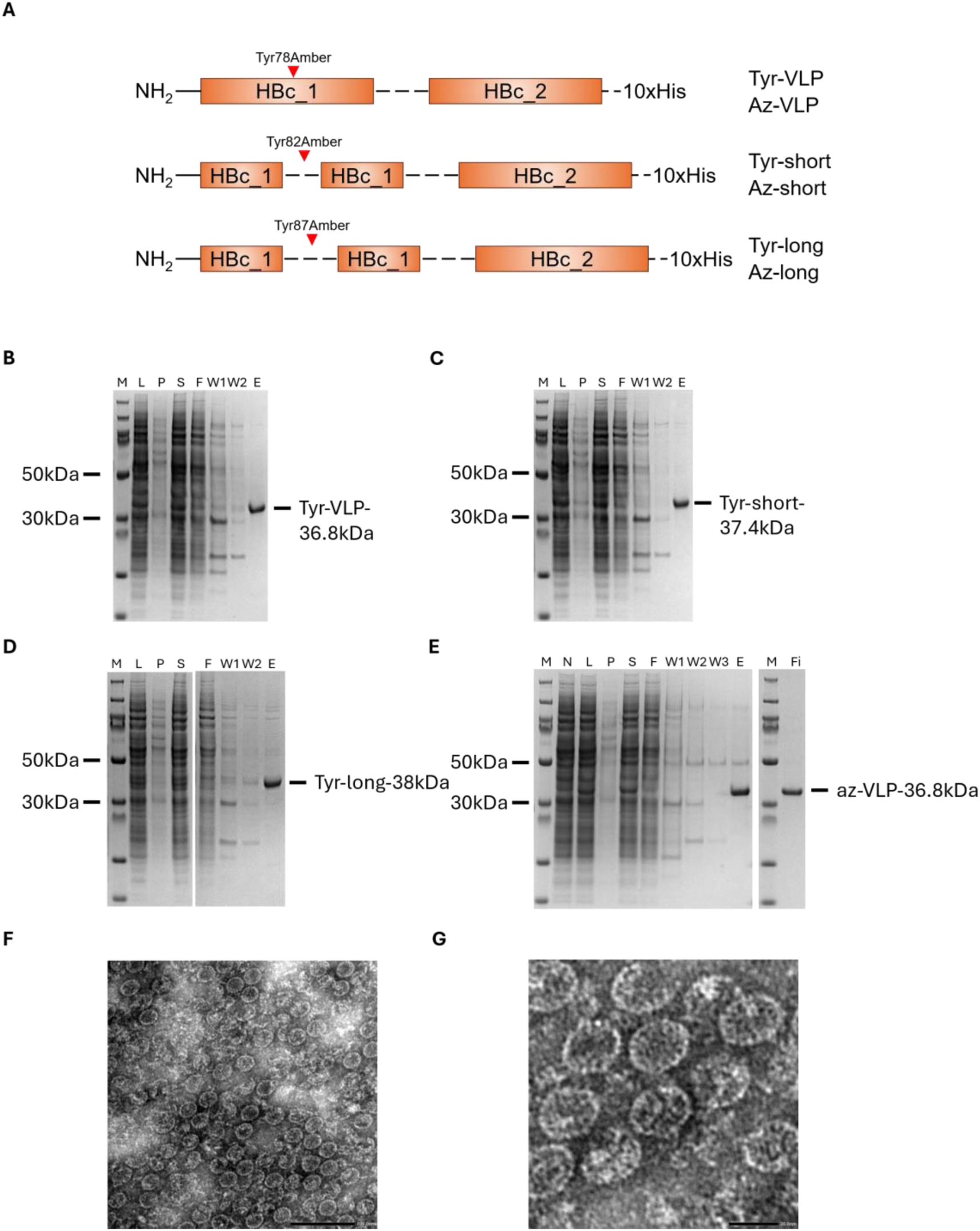
Expression and testing of different HBc VLP tandem core constructs in BYL. (A) Schematic representation of the VLP expression constructs used in this study. SDS-PAGE results showing the expression and IMAC purification of the (B) HBc VLP tandem core without linker (tyr-VLP), (C) HBc VLP tandem core with a short linker (tyr-short), (D) HBc VLP tandem core with a long linker (tyr-long) and (E) HBc VLP tandem core without linker containing the azido-tyrosine (az-VLP). (F, G) Transmission electron microscopy images for the az-VLP. The black bar indicates 100 nm and 20 nm, respectively. Abbreviations: M: molecular weight marker, L: lysate, P: pellet, S: supernatant, F: Flow-through, W: Wash, E: elution, Fi: Final sample, No: no linker HBc VLP; Short: short-linker HBc VLP; Long: long-linker HBc VLP; aVLP: azido-VLP without linker; N: non-template control; IMAC: immobilized metal ion chromatography. Created with BioRender.com.

The receptor binding domain (RBD) from influenza hemagglutinin was chosen as the antigenic partner to be conjugated to the azido-containing VLPs, achieved by expression in BYL and incorporation of alkyne-tyrosine. Four different hemagglutinin RBD protein constructs were designed with either the tyrosine (tyr-RBD) or ncaa insertion site (al-RBD) at N-terminus or C-terminus (RBD-tyr & RBD-al) (Figure 6A). RBD constructs with the tyrosine insertion expressed in BYL was found in both the pellet and supernatant fractions after centrifugation, indicating insolubility (Figure 6, B&D). Therefore, a purification procedure including an in-column refolding step was established. The al-RBD construct showed 2x greater yields than the RBD-al construct and expressed in a single band (Figure 6, C&E). The RBD-al construct showed expression in two bands where the smaller band is suggested as a premature translation termination product, where the amber stop codon was not efficiently supplied with alkyne-tyrosine in time. However, this truncated RBD product did not co-purify as it lacked the C-terminal purification tag. All following experiments were performed with the N-terminally tagged RBD construct given its higher yields.

**Figure 6.**
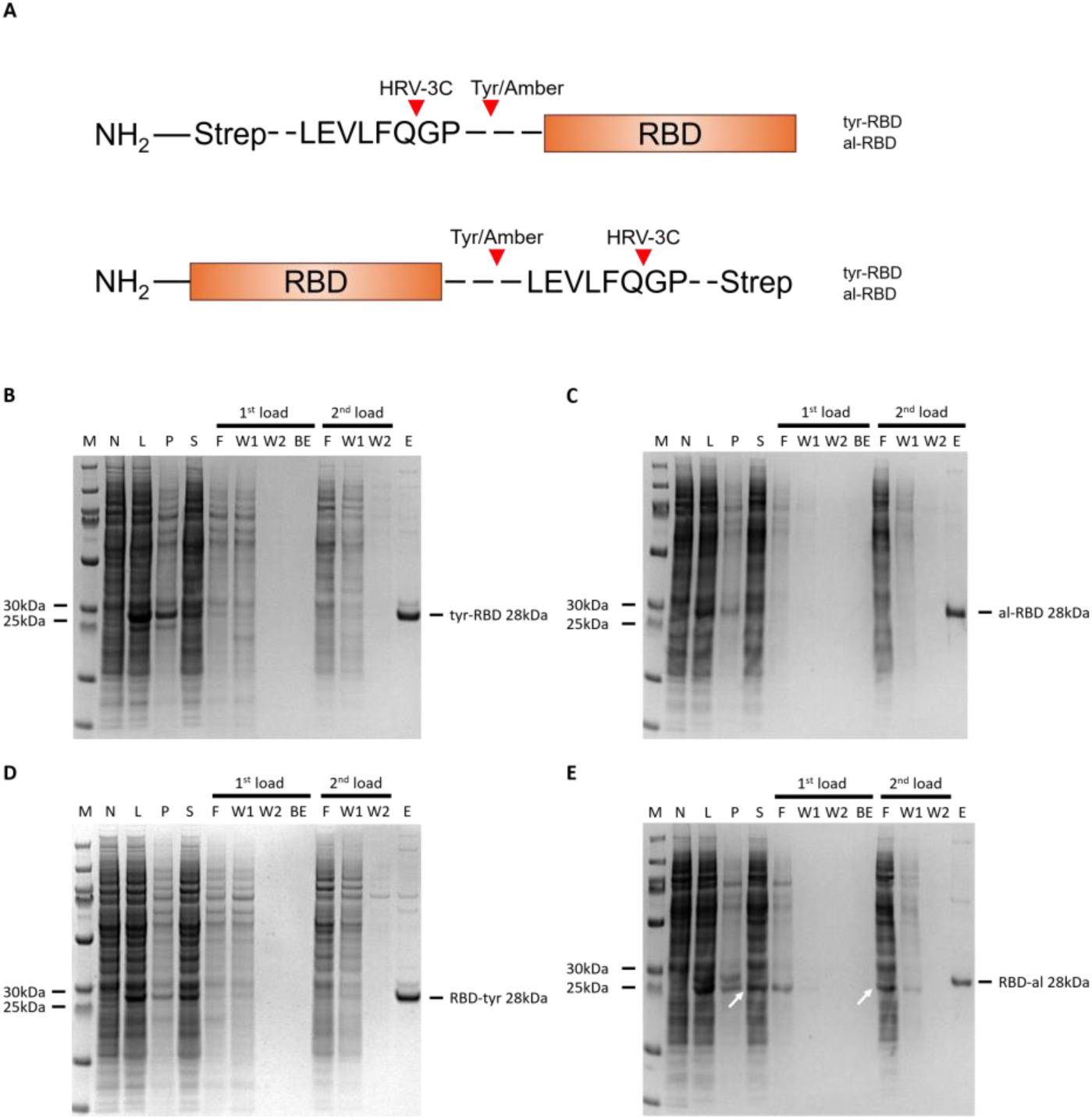
Hemagglutinin RBD expression and purification in BYL. (A) Schematic representation of the RBD expression constructs used in this study. SDS-PAGE results showing the expression and strep tag-mediated affinity purification of the different RBD constructs (B) N-terminally tagged tyr RBD (C) N-terminally tagged alkyne RBD (D) C-terminally tagged tyr RBD and (E) C-terminally tagged alkyne RBD. The purification process consisted of the solubilization and in-column re-folding of the RBD found in the pellet fraction (1^st^ load), and subsequent loading of the soluble BYL fraction (2^nd^ load). White arrows indicate truncated RBD without strep tag produced as failure of the amber suppression. Abbreviations: M: molecular weight marker, N: non-template control, L: lysate, P: pellet, S: supernatant, F: Flow-through, W: Wash, E: elution, BE: buffer exchange; RBD: receptor-binding domain. Created with BioRender.com.

### 3.5 Development of a plug-and-play VLP vaccination platform via click chemistry

CuAAC click chemistry reactions were utilized to drive the conjugation between az-VLPs and al-RBDs produced in BYL, thus also allowing to test the activity of the introduced ncaas. CuAAC reactions in the presence of tyr-VLP were optimized utilizing the 3-azido-7-hydroxycumarin (3a7) fluorescent dye. 3a7 was selected for its fluorogenic ‘click’ labelling properties, as it only becomes fluorescent upon successful CuAAC conjugation, thus giving a quantifiable response upon reaction completion. Higher Cu concentrations and the inclusion of Ni were required for the CuAAC reaction to take place in presence of HBc VLP (Figure 7A). These conditions were then used in the binding reactions between the az-VLP and al-RBD. As it can be observed, conjugation reactions took place as expected, showing the formation of the conjugated HBc-RBD product (Figure 7B). Modulating the concentration of each partner allowed the regulation of conjugation yields, from 30% to 70% (Figure 7C). Diminishing returns in conjugation yield were observed upon the addition of higher molarities of RBD. SEC analysis of RBD-VLP conjugates showed that most conjugate exists within the VLP peak, and a very little proportion as free conjugate (Figure 7D).

**Figure 7.**
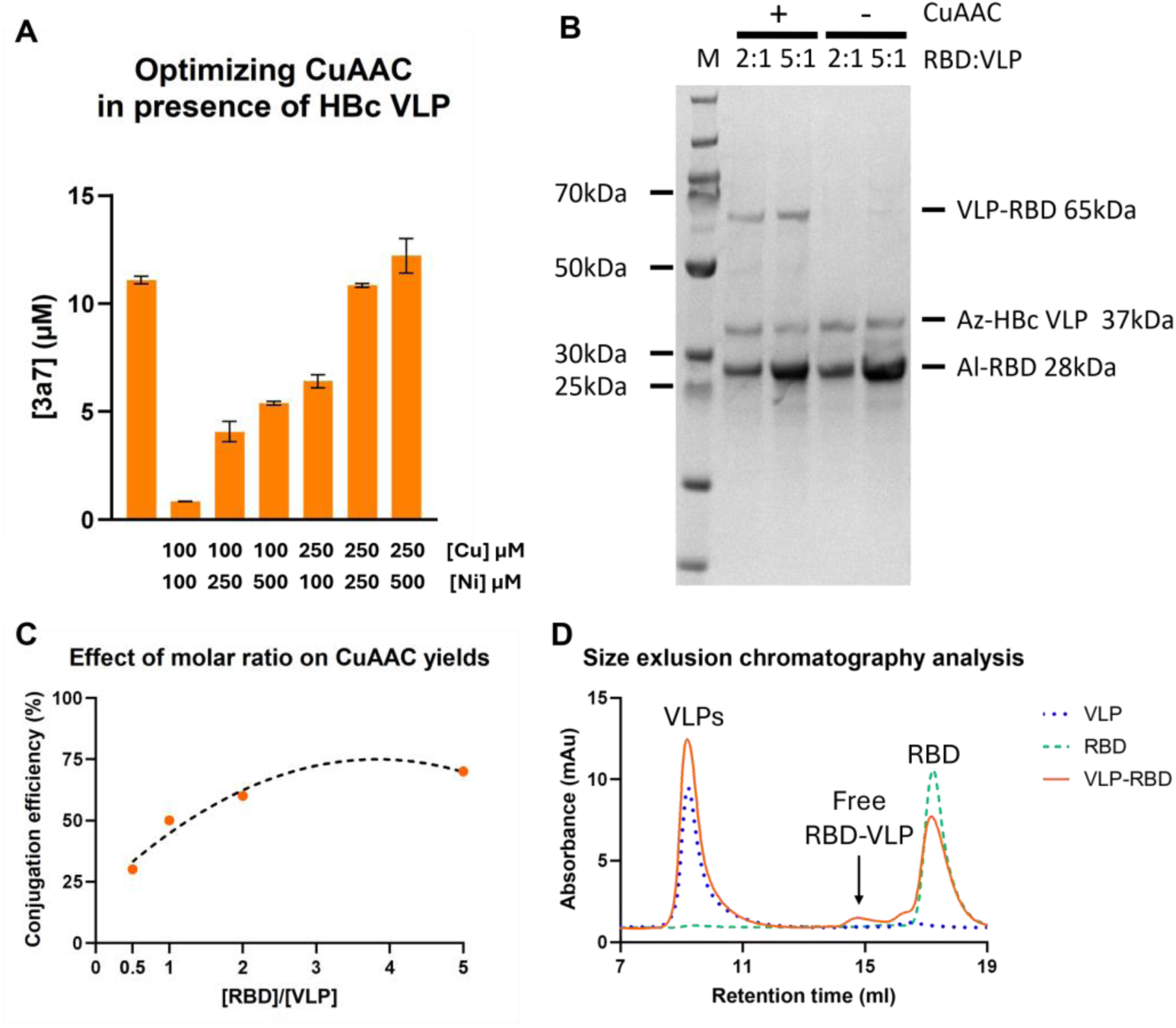
HBc VLP and hemagglutinin RBD CuAAC conjugation. (A) Fluorescence intensity analysis of CuAAC reaction of 3a7 and propargyl alcohol in presence of HBc VLP, performed under the indicated reaction conditions. (B) SDS-PAGE results of the CuAAC reactions between RBD and HBc VLP at the indicated molar ratios. The negative controls (-) indicate samples which did not undergo CuAAC. (C) Effect of modulating molar ratio between RBD and VLP on the conjugation yields. The % of RBD-VLP conjugate against free VLP is indicated. (D) Size-exclusion chromatography of the indicated samples. Abbreviations: 3a7: 3-azido-7-hydroxycoumarin; SEC: size-exclusion chromatography; VLP: virus-like particle; RBD: receptor binding domain.

Subsequently, free RBD was removed from VLP-RBD conjugates via SEC (Figure 8A). TEM analysis of the VLP-RBD conjugates showed that the VLP structure remained intact, whilst VLPs acquired a spiked form when compared to the non-conjugated VLPs, indicating the conjugation of RBD to their surface (Figure 8A & Figure S7). To test the sialic-acid binding functionality of the RBD-VLP conjugates, their ability to hemagglutinate chicken erythrocytes was tested. Free RBD and VLP, either on their own or combined without conjugation, did not lead to hemagglutination, whilst VLP-RBD conjugates could induce hemagglutination at concentrations higher than 1.25 ng/ml (Figure 8B). This indicates both, the sialic acid binding activity of the RBD proteins, together with their proper high avidity presentation on the VLP surface.

**Figure 8.**
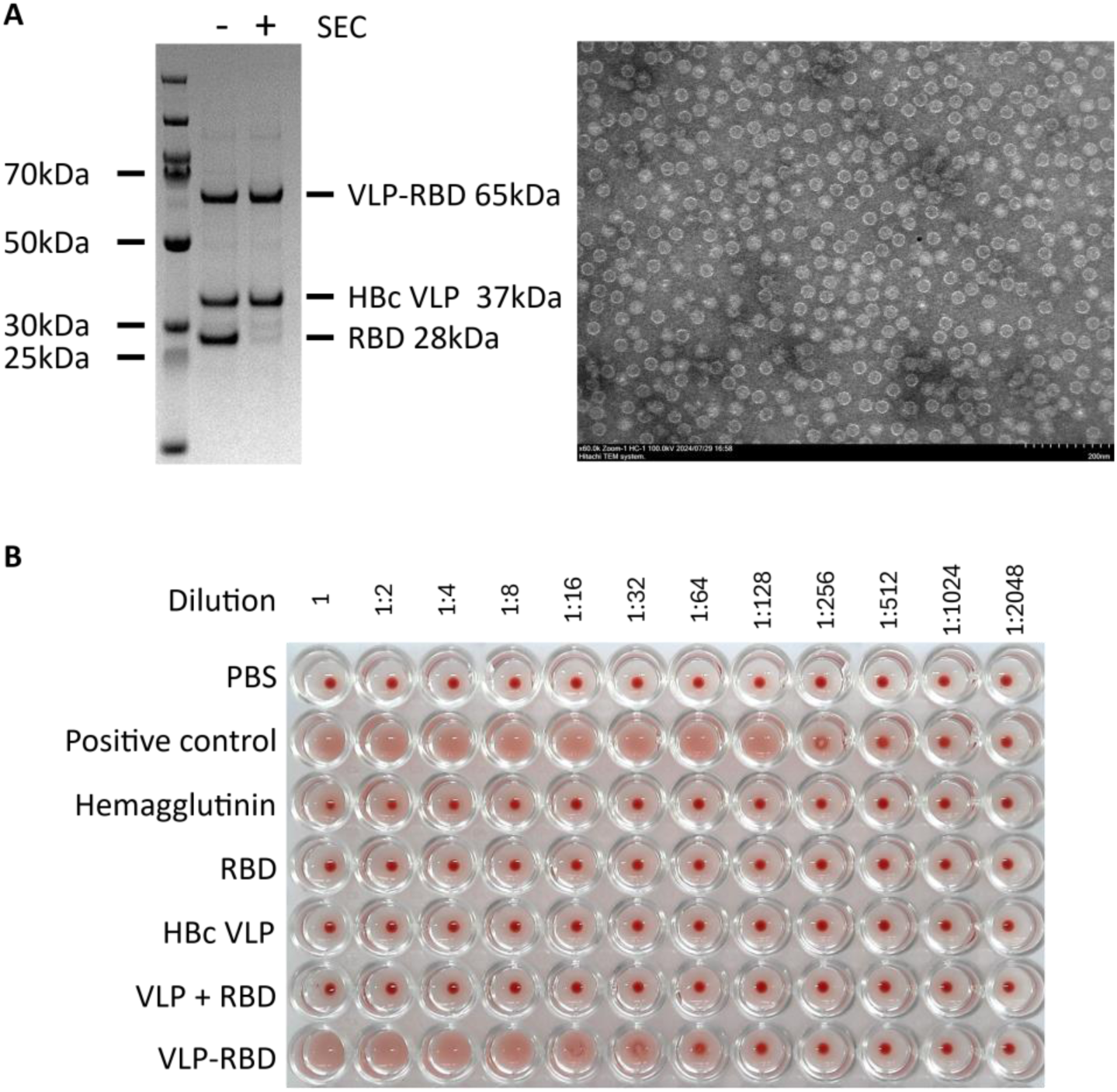
Analysis of VLP-RBD conjugates. (A) Coomassie blue-stained SDS-PAGE gel showing the removal of free RBD from VLP-RBD conjugates (left) and TEM image from the resulting conjugates (right, scale bar 200 nm). (B) Hemagglutination assay showing sialic-acid binding properties of the VLP-RBD conjugates. For tested samples, 4 µg of protein were plated in the initial column, and a serial dilution was performed as indicated. Commercial inactivated influenza vaccine (positive control), and hemagglutinin trimer protein (Hemagglutinin) were used as controls. Abbreviations: SEC: size-exclusion chromatography; HBc VLP: hepatitis B-core VLP; RBD: receptor binding domain from influenza hemagglutinin.

### 3.6 Protective efficacy of ncaa-enabled VLP–RBD conjugates in an influenza challenge model

Next, we wanted to test whether the immune response induced by BYL produced hemagglutinin antigens was protective against influenza infection, and whether any differences would be observed regarding conjugation to the HBc VLP. For that purpose, mice were immunised with one of the following formulations: free RBD, RBD conjugated to VLP, free RBD combined with free VLP, and free VLP as negative control. Mice were immunised by the intramuscular route in a prime-boost regime with a 5 µg dose of each formulation (Figure 9A). Mice were then challenged intranasally with a mouse-specific influenza strain: PR/8 pH1N1. Viral load after infection was lower in the RBD-vaccinated mice, without any significant difference between VLP-conjugated or un-conjugated RBD (Figure 9B). RBD-VLP conjugated vaccinated mice did not lose weight after infection; whilst the RBD alone and RBD-VLP unconjugated were partially protected (Figure 9C). After one dose of vaccine, mice immunised with RBD or RBD conjugated to VLP had higher titres of anti-PR/8 IgG than the unconjugated or the VLP alone groups (Figure 9D). After 2 doses of the antigen all three groups receiving some RBD had higher titre than the VLP alone group (Figure 9D); with the RBD-VLP conjugated group showing the highest value. These results show the potential of BYL to produce protective antigens against influenza infection and that multivalent presentation from ncaa-mediated conjugation to VLPs can boost the protective effect against disease Altogether, the introduction of ncaas into the BYL CFPS system was enabled, without impacting protein yields whilst retaining its scalability. We could apply ncaa incorporation to generate a plug-and-play vaccine platform where the HBc VLP could be conjugated to extrinsic antigens such as hemagglutinin RBD, not only retaining the structure and functionalities of both elements but boosting their protecting qualities.

**Figure 9.**
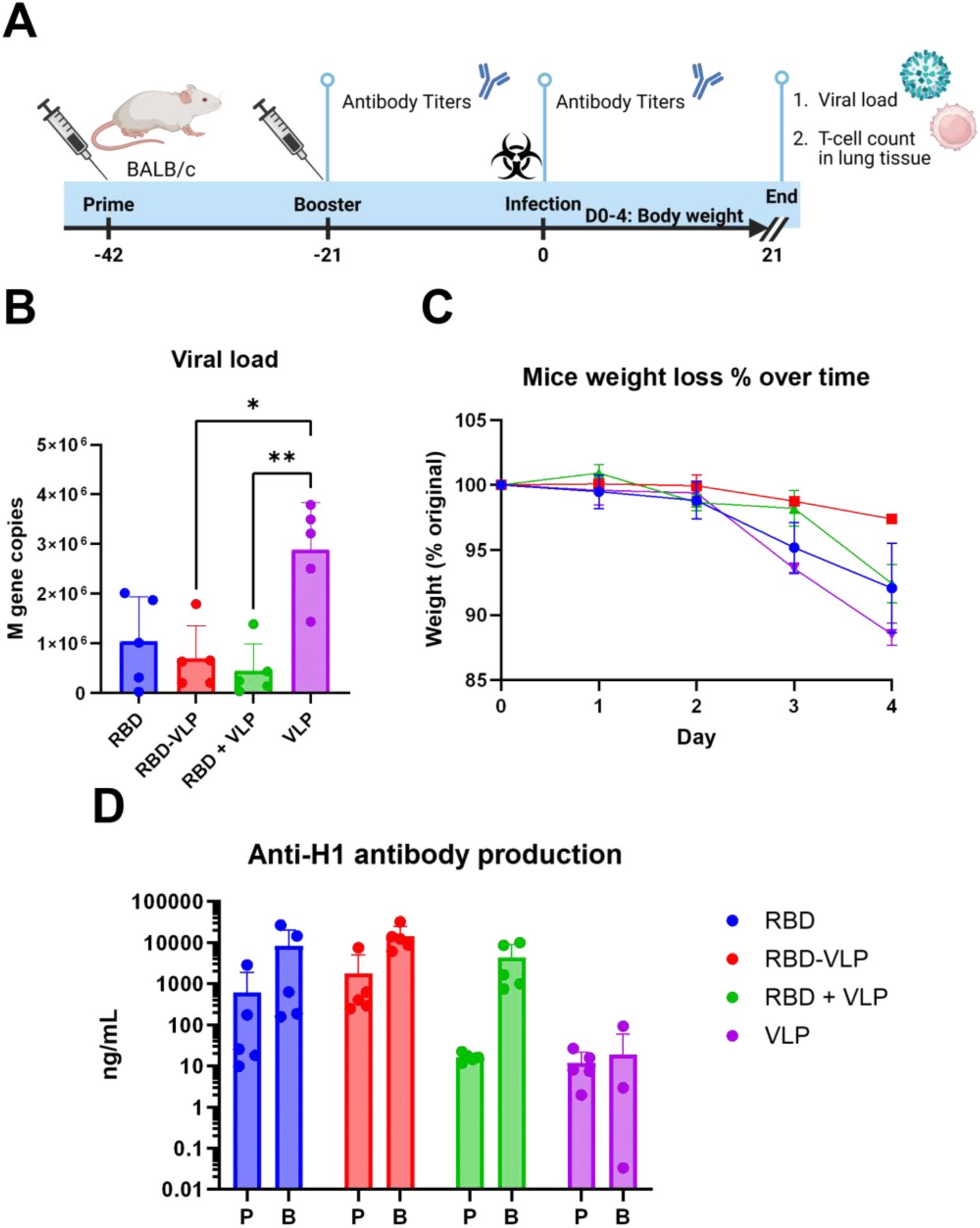
Mouse PR8 influenza challenge study. **(A)** Mice were immunised two times at 3 weekly intervals with intramuscular admission of 5μg RBD for each indicated vaccine formulation. Three weeks after the first and second immunisations, anti-HA antibody titre was measured by ELISA. Mice were then challenged with 3x10^3^ PFU PR/8 by the intranasal route. **(B)** Viral load on day 4 after infection measured by RT-PCR A. **(C)** Weight change was assessed after infection. **(D)** Anti-HA antibody titre measured by ELISA after prime (P) and Booster (B). Points represent individual animals (B, D) or mean (C) of n=5 mice per group. Abbreviations: P: primer; B: Booster; RBD: receptor-binding domain from influenza hemagglutinin; VLP: HBc Virus-like particle.

## 4. Discussion

In this study, we successfully implemented the introduction of non-canonical amino acids (ncaas) into proteins produced in the BYL CFPS system commercialised as ALiCE®. The *E. coli* TyrT amber suppression system showed high incorporation yields for both, azido and alkyne tyrosine ncaas, with up to 100% of incorporation efficiency when purified transferase was supplemented to BYL reactions. We then enabled the linear scaling of ncaa incorporation in BYL by uncoupling tRNA transcription from CFPS reactions. Ncaa incorporation in BYL was applied to vaccine development by the post-assembly conjugation of HBc VLPs and influenza RBD proteins via CuAAC-mediated click chemistry. Conjugation yields could be easily regulated by modifying the concentration of the respective partners and VLPs retained their structure after conjugation. VLP-RBD conjugates could efficiently bind sialic acid in a hemagglutination assay and showed potential in increasing the protective effect of VLP-conjugated antigens *in vivo*. Together, these results show that the high-yielding BYL CFPS system can be used for the efficient introduction of ncaa into recombinant proteins.

Given the relevance of CFPS systems, especially in their ability to introduce ncaa into recombinant proteins, we aimed to establish ncaa-incorporation into the eukaryotic BYL CFPS system (10,49,50). For this purpose, we studied the *E. coli* tyrosyl (eTyrT), and pyrrolysine *M. mazei* (mPyrT) and *M. barkeri* (bPyrT) amber suppression systems. All these systems utilize a combination of matching amber-supressing tRNA, aminoacyl-transferase, and ncaa. For maximally efficient ncaa incorporation, it was essential to optimize the formulation and format of these different elements for use in BYL. For instance, the solubility problems of the pyrrolysine transferases tested in this project are known and have been addressed in literature in a myriad of ways (41–44). In our studies, fusion with Smbp solubility tag did not notably increase transferase solubility, in contrast to what was previously observed in *E. coli* expression (45). The eTyrT, on the other hand, could be efficiently expressed and purified, allowing it to reach the higher concentrations required for efficient aminoacyl-transferase activity within the BYL CFPS reaction.

The efficient transcription of the amber suppressor tRNA with sequence fidelity is essential for recognition and aminoacylation by the transferase (46). In our studies, we observed that the design of the tRNA and format of the corresponding DNA template were essential factors for ncaa incorporation. tRNA constructs with hammerhead ribozyme sequences showed lower ncaa incorporation yields with the *E. coli* tyrosine transferase but improved yields with the pyrrolysine transferases, thus indicating different ribozyme tRNA processing behaviours, as observed with the presence of bigger tRNA species. However, the biggest impact on ncaa incorporation efficiency came from using oligonucleotides and linearized plasmid for tRNA, where oligonucleotide-mediated reduction in yield is theorized to originate from an accumulation of synthesis errors for these long primers (ca. 54% fidelity according to manufacturers (47)). The yield decrease when utilizing linearized plasmid could be explained by the absence of PTO and 2-O-methoxy modifications, that would prevent exonuclease reaction within BYL (48), and untemplated 3’ transcription (30), respectively. We took these insights into consideration for the scaling of ncaa incorporation in BYL, summarized in **Table 1**, where the different estimated cost effectiveness of each tRNA approach, together with their yields and simplicity are subjectively rated. As it can be observed, even if the *in vitro* transcription of the PCR template is not the simplest of approaches, it provides the highest ncaa incorporation yields, whilst maintaining lower costs and showing linear scalability. Further improvements could be performed to the process, including the *in vitro* aminoacylation of the tRNA transcript, which could in turn further reduce tRNA, amino acyltransferase and ncaa requirements (49).

**Table 1.**
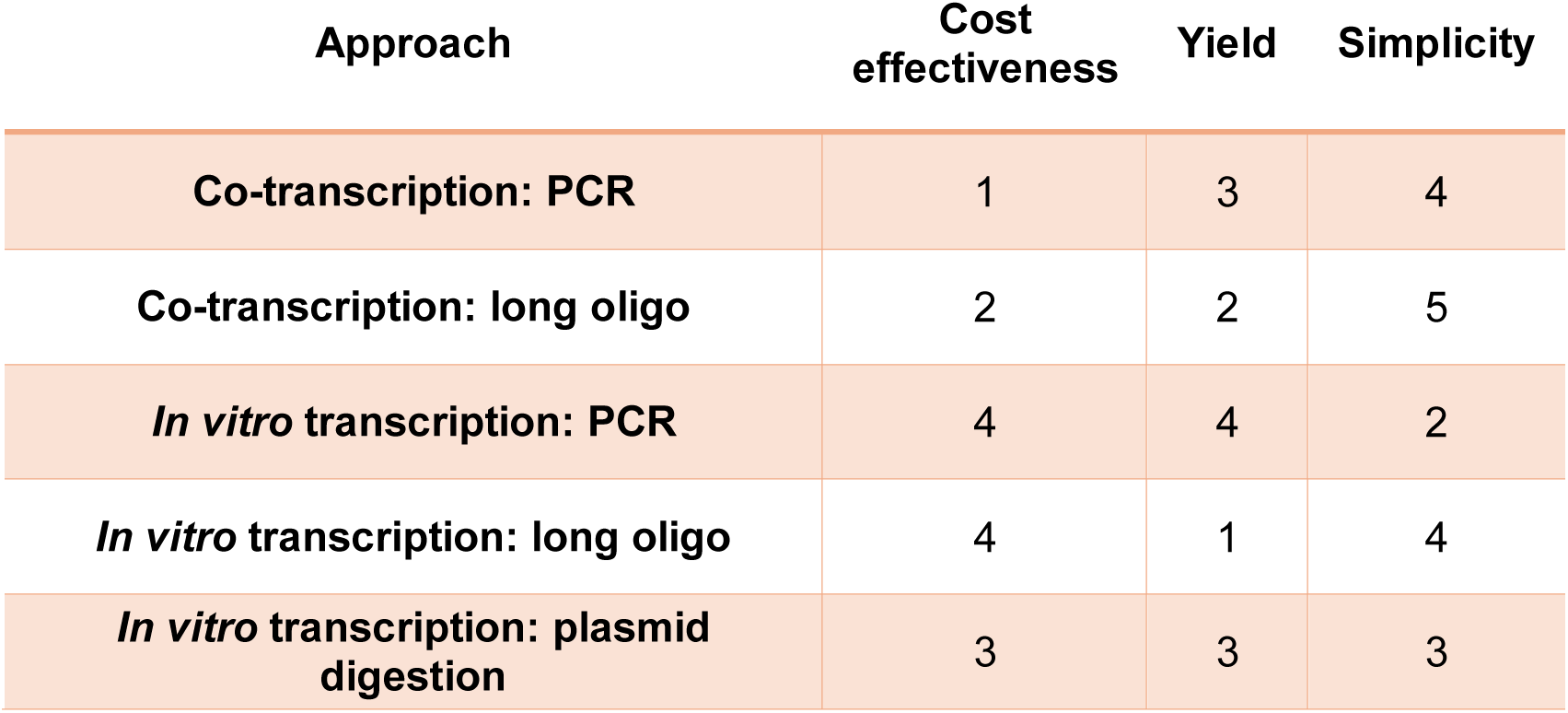
Cost estimation for the different tRNA templates and approaches. A subjective summary statistic for the yield, and simplicity of each approach is also provided, with a 1-5 scoring scale, where 1 is the lowest and 5 the highest.

Finally, the manner all ncaa incorporation components are integrated is also of critical importance. We observed that supplementation of unpurified transferase from a running reaction led to a considerable reduction in overall BYL protein yields, caused by the competition over the transcription and translation machinery between the transferase and the a-eYFP construct (50). This was addressed via transferase purification and supplementation, showing great azido and alkyne-tyrosine incorporation efficiencies when using the eTyrT system, whilst the mPyrT still showed very low incorporation yields. Optimizations might still be possible, including the tRNA template, aminoacyl-transferase and ncaa concentration, or altering codon usage around the amber site to decrease translation speed (51). Given that only a single release factor is responsible for translation termination in eukaryotes, its silencing as performed in *E. coli* would not be possible (52).

Subsequently, we aimed to apply ncaa incorporation in BYL to vaccine development. For that purpose, we used ncaa-mediated click-chemistry to combine the enhanced immunogenic properties of the HBc VLP, with the known influenza vaccine antigen, hemagglutinin. The HBc VLP is a well characterized nanoparticle with known assembly mechanisms and previously described applications as a carrier VLP for this purpose, also for drug delivery (53). We have shown the high yield and scalable production of HBc VLPs in BYL (32), and in this research we have furthered its applicability for vaccine production by introducing conjugation handles. The RBD of the hemagglutinin protein was chosen as a conjugation partner for the HBc VLP, given its proven capabilities to fold into its native form and triggering protective immune responses in mice (54,55). RBD-VLP conjugates showed hemagglutination activity, thus proving the sialic-acid binding activity of the RBD proteins, and their proper high-avidity presentation on the surface of the VLPs.

When used to vaccinate mice, conjugation of RBD to VLP enhanced the protection against the development of disease, observed as lower weight loss after homologous viral challenge. Previously, SpyTag/SpyCatcher conjugation to synthetic VLPs was utilized to increase the immunogenicity of SARS-CoV-2 RBD proteins (56). Conjugating RBD to VLPs significantly boosted antibody levels and ACE2-blocking activity in immunized mice, even at a low dose of 5 µg, compared to free RBD. In contrast, standalone SARS-CoV-2 RBD can induce neutralizing antibodies on its own but typically requires much higher doses (50–100 µg) (57). Similarly, in the case of *E. coli*-produced influenza hemagglutinin RBD, relatively high vaccine doses ranging from 125 to 200 µg were needed to achieve a protective immune response in ferrets (58). However, we found that unconjugated influenza RBD could also provide a considerable immune response in mice, possibly explaining why we did not observe a significant difference between free-RBD and VLP-conjugated RBD on antibody titre or viral load. Nonetheless, this data serves as a first proof of concept for the potential of VLP conjugation to increase the immunogenicity and/or protection properties of a given vaccine candidate. Future research aims to study the effect of different dosages, as well as modulating the degree of VLP-antigen conjugation to optimize the protective response. Moreover, this same conjugation approach could be utilized to conjugate antigens from different pathogens to the HBc VLP, thus resulting in a plug-and play vaccine platform applicable to pandemic preparedness, whereby the adjuvating carrier VLP can be stockpiled in advance needing only to produce the necessary antigens.

All in all, this study shows the promising possibilities to use the BYL CFPS system to produce proteins containing non-canonical amino acids in high yields. The benefits of the BYL system regarding its scalability, yields and production of complex proteins, together with the added properties of non-canonical amino acids, further increase the potential of BYL for the screening and production of proteins of clinical relevance, such as vaccines and antibody-drug conjugates.

## Acknowledgments

We thank Eva Miriam Buhl (Uniklinikum Aachen) for her assistance with the transmission electron microscopy experiments and Jonathan Fauerbach for his assistance in reviewing this manuscript.

