## Supplemental figures for "Le click c’est chic: a plug-and-play virus-like particle vaccination platform enabled by non-canonical amino acid incorporation and click chemistry in the tobacco BY-2 cell-free protein synthesis system"

Supplemental data

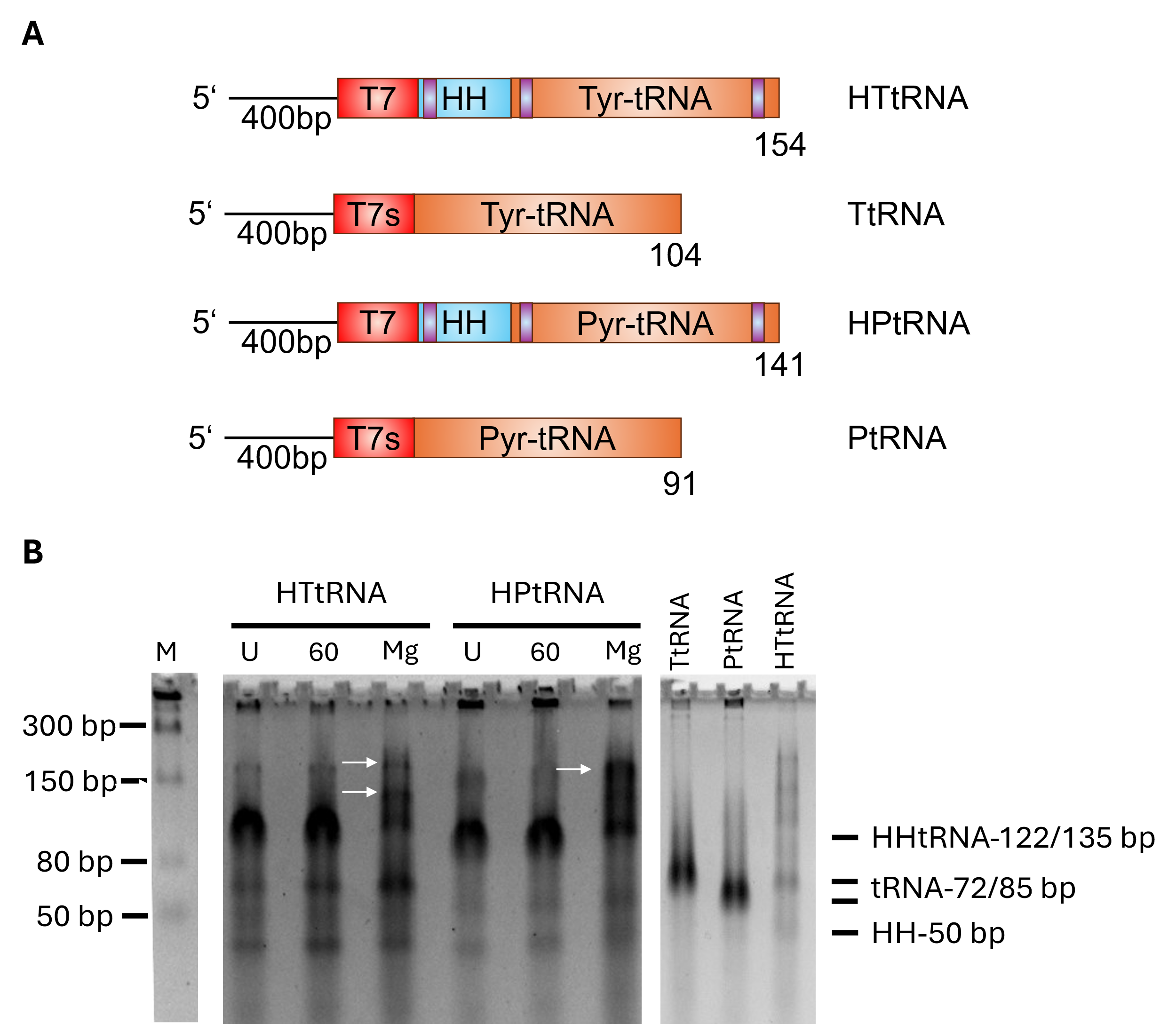

**Figure S1 – tRNA construct design and transcription. (A)** Graphical representation of the tRNA constructs used in this study. Constructs either containing a self-cleaving hammerhead ribozyme or shorter T7 promoter were designed as to ensure 5’ homogeneity. The hybridization box for the hammerhead ribozyme is marked in purple. Note that the tRNA used for mPyrT and bPyrT is the same. **(B)** IVT and agarose gel electrophoresis of tRNA constructs. Left gel shows the transcription from constructs containing a hammerhead sequence. U indicates the untreated tRNA directly after IVT, 60 the tRNA after 1h 60°C treatment and Mg samples treated for 1h at 60°C with extra Mg added. White arrows indicate cleavage products of greater size resulting from the hammerhead activity. White arrows could indicate the formation of multimeric forms upon cleavage. Right gel shows the transcription of the hammerhead-less constructs, with one hammerhead construct after 60°C+Mg treatment as comparison. Abbreviations: U: untreated; T7: T7 promoter; T7s: shorter T7 promoter, HH: hammerhead ribozyme; M: molecular weight marker. Created with BioRender.com.

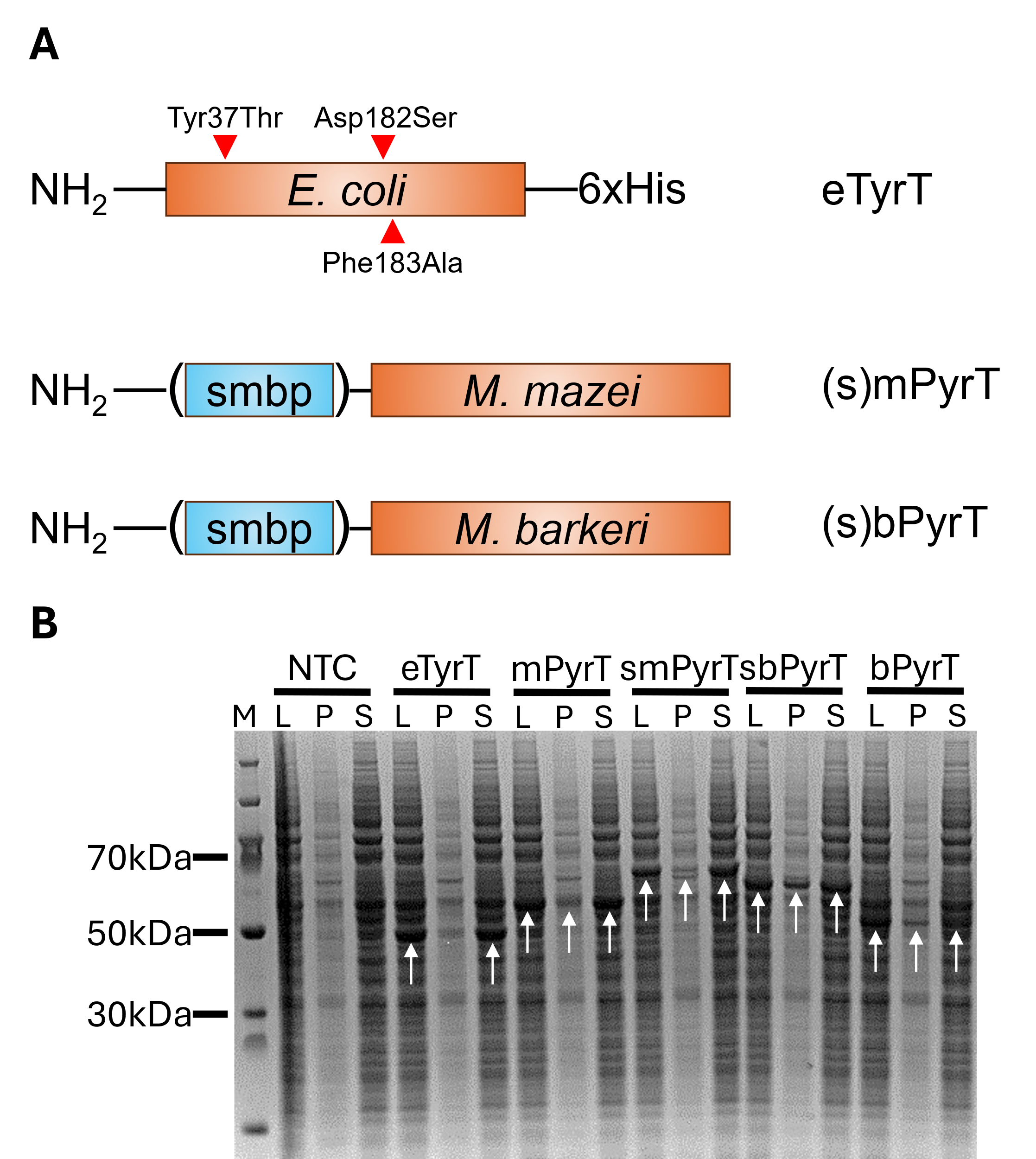

**Figure S2 - Testing BYL expression of the different amber suppression systems.** (A) Graphical representation of the aminoacyl-transferase expression constructs used in this study. For *M. mazei* and *M. barkeri*, each construct was generated with and without a smbp tag to increase solubility, resulting in four variants. (B) Coomassie blue-stained SDS-PAGE results showing the BYL expression of the different aminoacyl-transferases. The expected size in kDa for each protein is: eTyrT: 47, mPyrT: 51, smPyrT: 61, sbPyrT: 57,5 and bPyrT: 47,5. White arrows indicate the protein band for each respective transferase. Abbreviations: smbp: small-metal binding protein; M: molecular weight marker; L: lysate; P: pellet; S: supernatant; NTC: non-template control; Smbp: small-metal binding protein; His: poly-histidine tag. Created with BioRender.com.

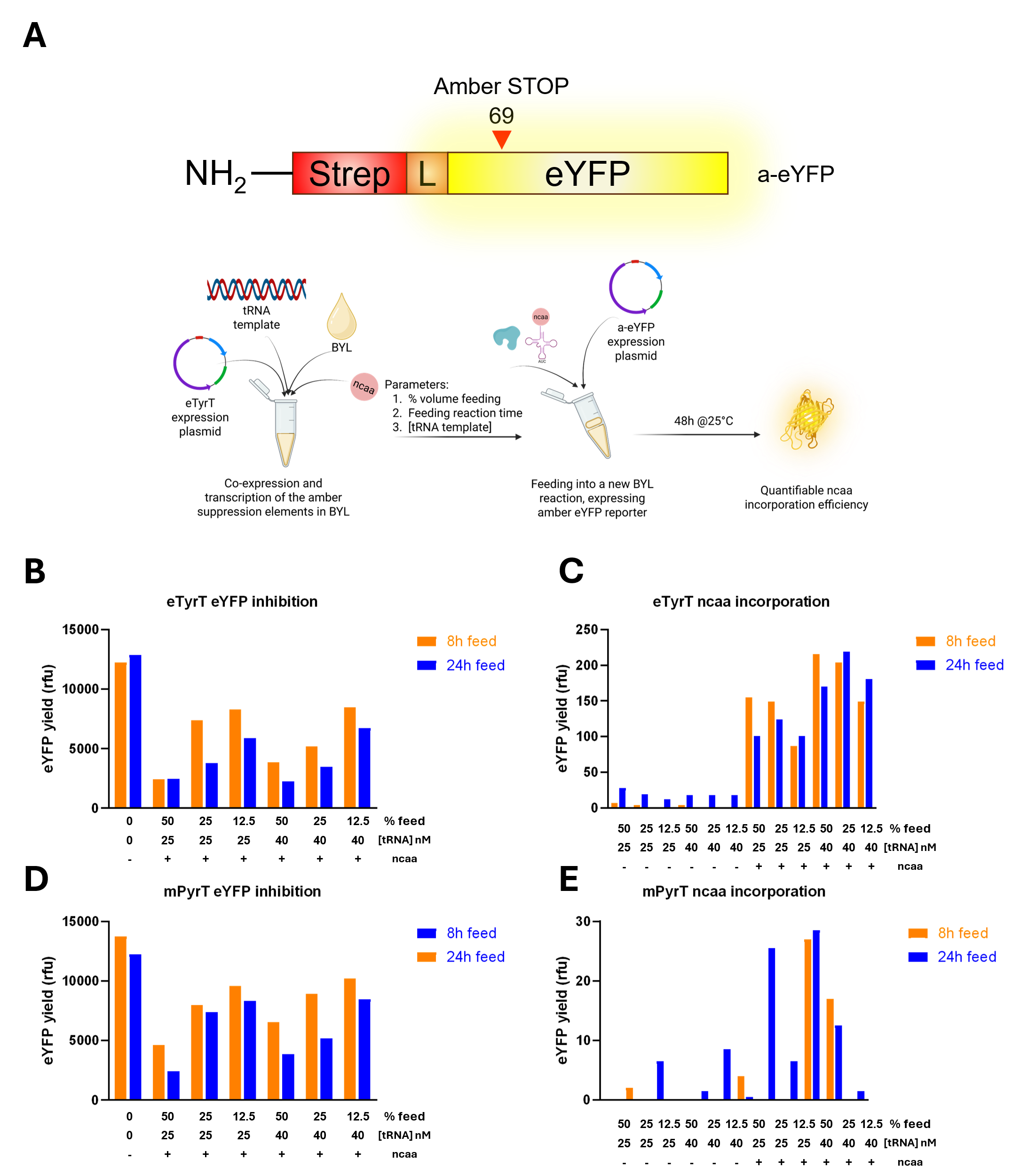

**Figure S3 - BYL ncaa incorporation efficiency via aminoacyl-transferase co-expression and supplementation**. (A) Schematic representation of the amber-eYFP expression construct and supplementation approach used in this study. Inhibitory effect on the BYL reaction when supplementation the indicated volumes of BYL expressions containing the eTyrT (B) and mPyrT (C) amber suppression components. Ncaa incorporation into a-eYFP when supplementation the indicated volumes of BYL expressions containing the eTyrT (D) and mPyrT (E) amber suppression components. Bars indicate measurements from single experiments. Abbreviations: ncaa: non-canonical amino acid; eYFP: enhanced yellow fluorescent protein; a-eYFP: amber-containing eYFP; RFU: relative fluorescence units. Created with BioRender.com.

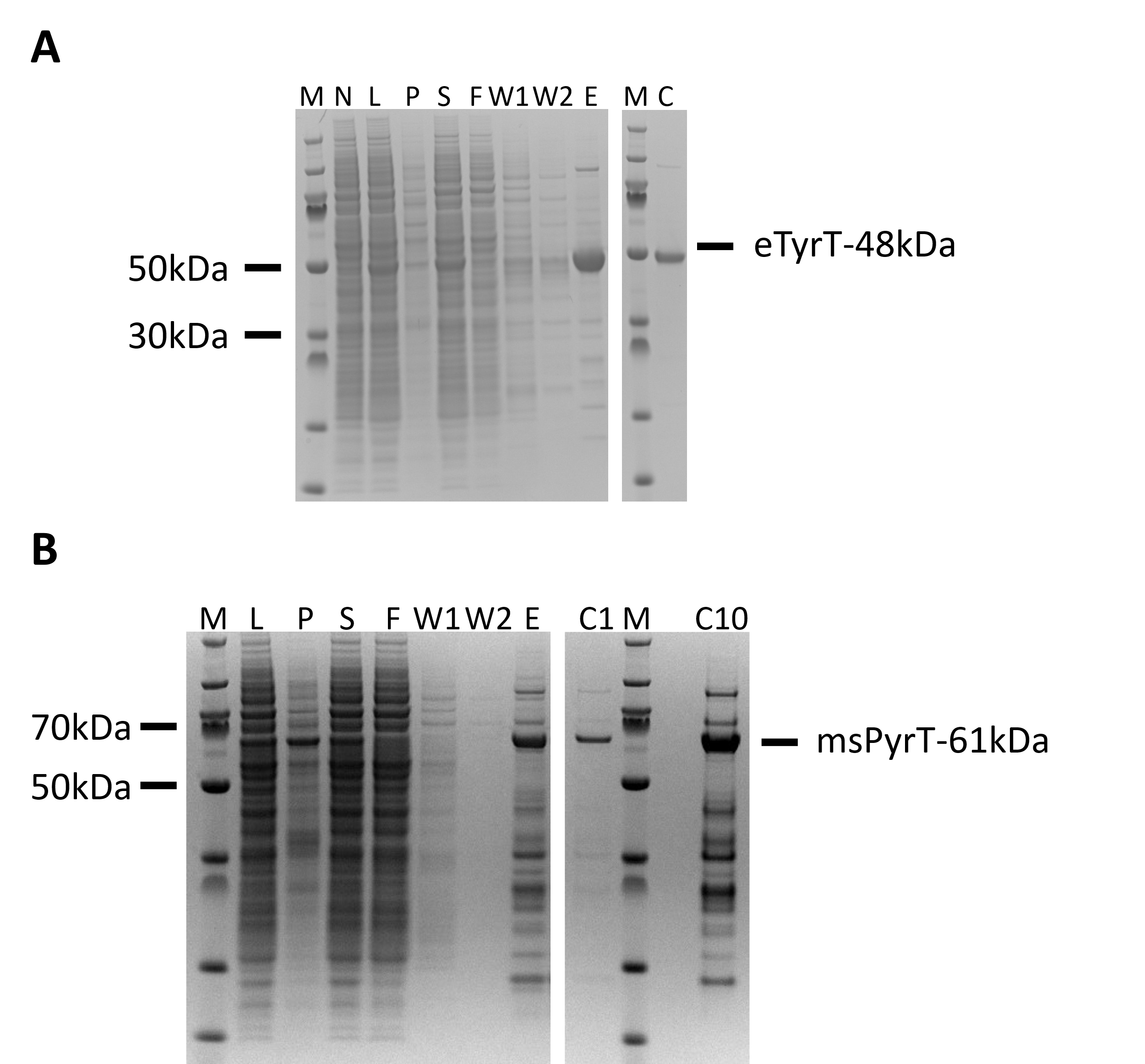

**Figure S4 - BYL expression and purification of aminoacyl-transferases.** Coomassie-blue stained SDS-PAGE gels showing the expression and IMAC purification of the eTyrT (A) and smPyrT (B) aminoacyl-transferases. Abbreviations: M: molecular weight marker; N: non-template control; L: lysate after BYL reaction; P: pellet after centrifugation; S: supernatant after centrifugation; F: flow through; W: wash; E: elution; C: Concentrated final transferase; C1: 1µg of the final concentrated transferase; C10: 10µg of the final concentrated transferase.

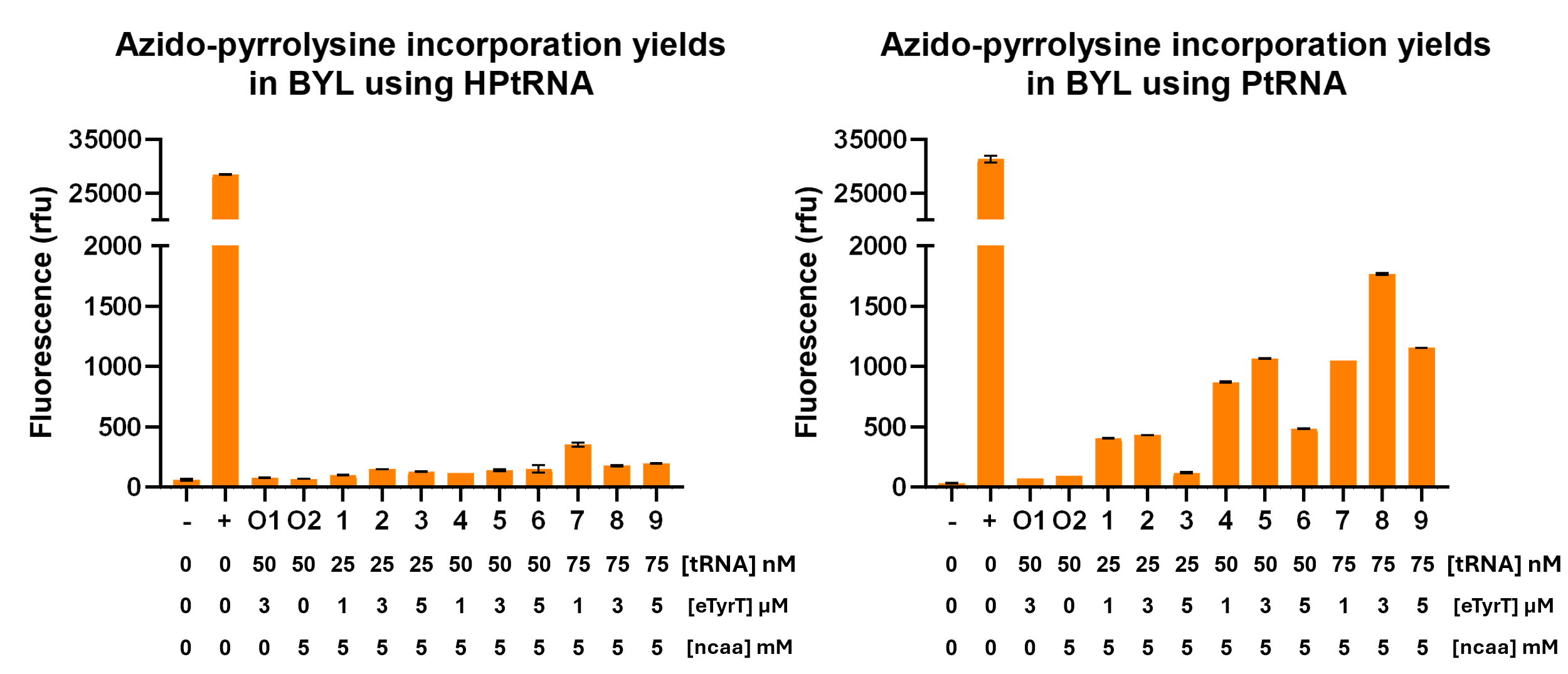

**Figure S5 - Ncaa incorporation efficiency of azido-pyrrolysine using mPyrT and different tRNA templates in BYL.** The incorporation efficiency was measured in terms of fluorescence of the amber eYFP construct after a 48h BYL reaction. Expression of eYFP is used as a positive control (indicated as +), indicating the maximum BYL yield, and the orthogonal controls 1 and 2 indicate the nonspecific incorporation of native amino acids and ncaa by native transferases, respectively. Different concentrations of the tRNA template and eTyrT enzyme were explored as indicated. Data points represent averages of experiments performed in duplicate. Abbreviations: rfu: relative fluorescence units; (-): Non-template BYL control; (+) pALiCE01 positive control; O1-2: orthogonality controls.

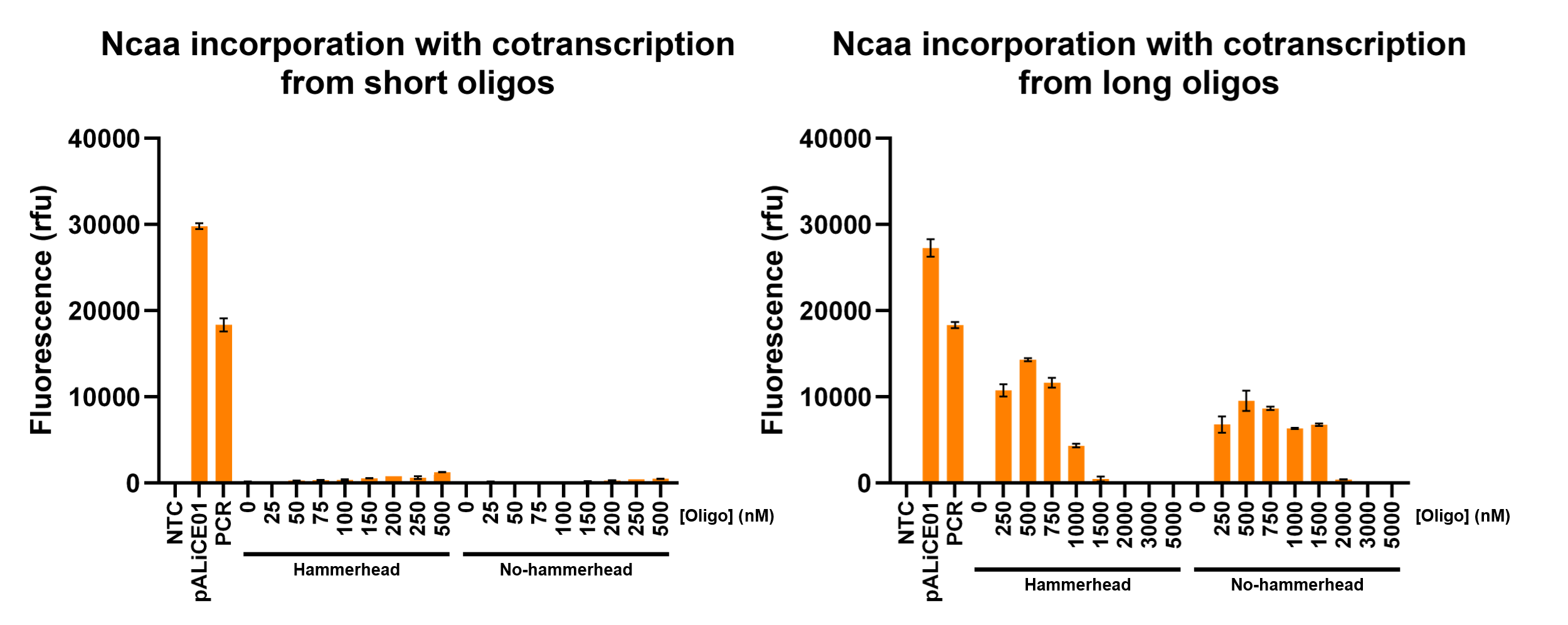

**Figure S6 – Ncaa incorporation in BYL when utilizing annealed oligos for co-transcription.** The incorporation efficiency was measured in terms of fluorescence of the amber eYFP construct after a 48h BYL reaction. Expression of eYFP from the pALiCE01 plasmid is used as a positive control, indicating the maximum BYL yield. The different concentrations of the annealed oligos in nM are shown. Data points represent averages of experiments performed in duplicate. Abbreviations: rfu: relative fluorescence units; (-): Non-template BYL control; (+) pALiCE01 positive control.

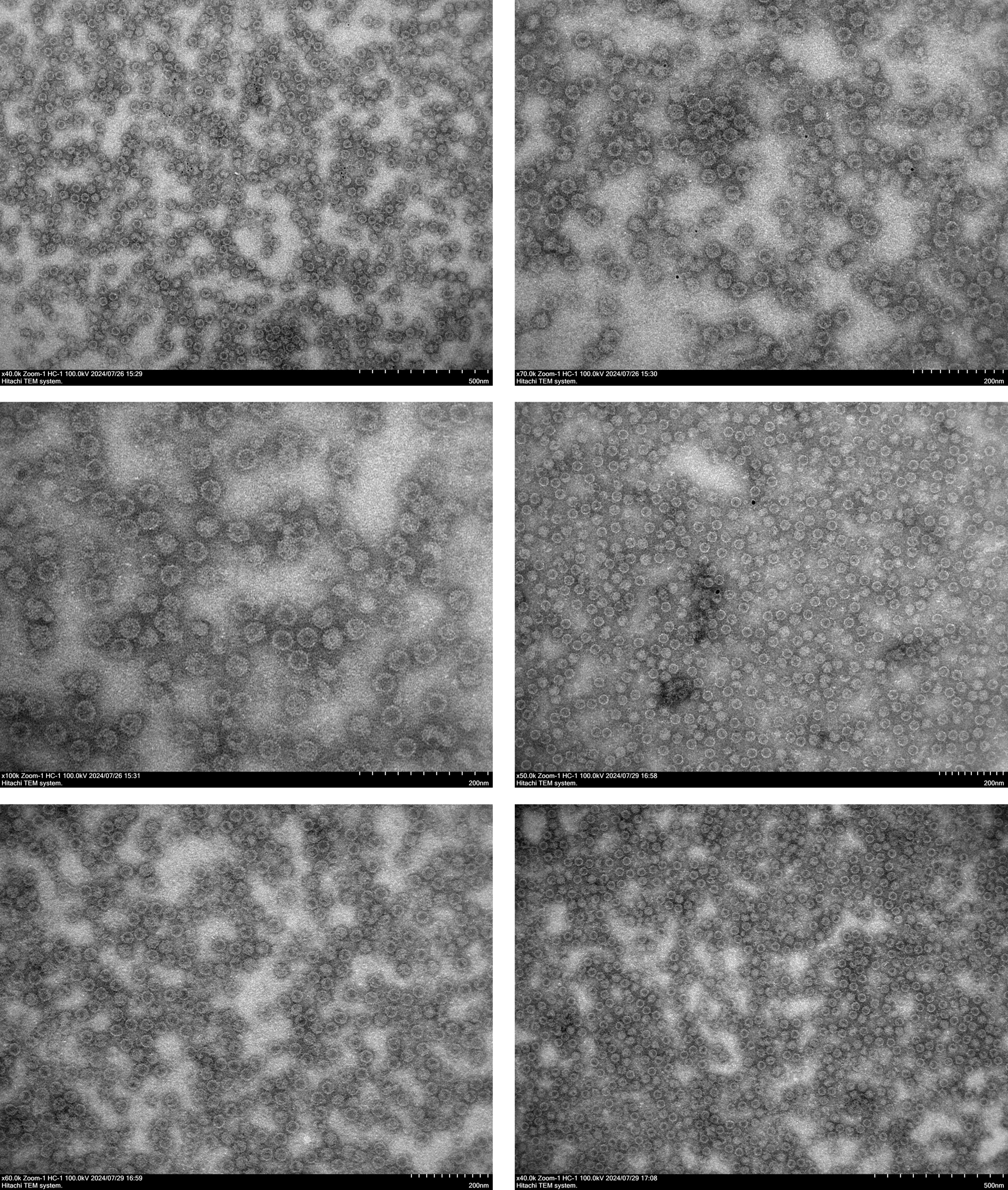

**Figure S7 - Transmission electron microscopy of HBc VLP-RBD conjugates.** The scale for each image is indicated in the bottom right (500 or 200 nm depending on the image). Abbreviations: HBc VLP: hepatitis B-core virus-like particle; RBD: receptor-binding domain of the influenza hemagglutinin.
